# The bacterial effector HopZ1a acetylates MKK7 to suppress plant immunity

**DOI:** 10.1101/2020.06.14.150508

**Authors:** José S. Rufián, Javier Rueda-Blanco, Diego López-Márquez, Alberto P. Macho, Carmen R. Beuzón, Javier Ruiz-Albert

## Abstract

The *Pseudomonas syringae* type III secretion system translocates effector proteins into the host cell cytosol, suppressing plant basal immunity triggered upon recognition of pathogen-associated molecular patterns (PAMPs), and effector-triggered immunity. Effector HopZ1a suppresses local and systemic immunity triggered by PAMPs and effectors, through target acetylation. HopZ1a has been shown to target several plant proteins, but none fully substantiates HopZ1a-associated immune suppression. Here, we investigate *Arabidopsis thaliana* mitogen-activated protein kinase kinases (MKKs) as potential targets, focusing on AtMKK7, a positive regulator of local and systemic immunity. We analyse HopZ1a interference with AtMKK7 by translocation of HopZ1a from bacteria inoculated into Arabidopsis expressing MKK7 from an inducible promoter. Reciprocal phenotypes are analysed on plants expressing a construct quenching MKK7 native expression. We analyse HopZ1a-MKK7 interaction by three independent methods, and the relevance of acetylation by *in vitro* kinase and *in planta* functional assays. We demonstrate AtMKK7 contribution to immune signalling showing MKK7-dependent flg22-induced ROS burst, MAPK activation, and callose accumulation, plus AvrRpt2-triggered MKK7-dependent signalling. Further, we demonstrate HopZ1a suppression of all MKK7-dependent responses, HopZ1a-MKK7 interaction in planta, and HopZ1a acetylation of MKK7 in a lysine required for full kinase activity. We demonstrate that HopZ1a targets AtMKK7 to suppress local and systemic plant immunity.

## INTRODUCTION

*Pseudomonas syringae* is a phytopathogenic bacterium that uses a type III secretion system (T3SS) to secrete proteins, known as effectors, directly inside the host cell cytosol. A number of *P. syringae* type III effectors (T3Es) can suppress the plant defence response triggered upon recognition by plant pattern recognition receptors (PRRs) of conserved pathogen-associated molecular patterns (PAMPs) such as bacterial flagellin, known as PAMP-triggered immunity (PTI) (Boller & Felix, 2009). In turn, T3Es can be detected, directly or indirectly, by plant resistance proteins containing nucleotide-binding domains and leucine-rich repeats (NLRs), triggering a stronger line of defense known as effector-triggered immunity (ETI), which usually ensues a type of programmed cell death referred to as the hypersensitive response (HR), resulting in a drastic restriction of pathogen growth (Chiang, 2015). Further, other T3Es can suppress ETI, thus enabling pathogen growth (Jones & Dangl, 2006). The local activation of immunity also triggers a plant defense response that goes beyond the local tissue, known as systemic acquired resistance (SAR) (reviewed by Spoel & Dong, 2012; Klessig *et al.*, 2018; Shine *et al.*, 2019).

HopZ1a is a *P. syringae* T3E with the ability to suppress plant immunity, including (*i*) basal resistance or PTI triggered by *P. syringae* pv. tomato (*Pto*) DC3000 (Macho *et al.*, 2010; Lewis *et al.*, 2014); (*ii*) ETI triggered by the heterologous effectors AvrRpt2, AvrRps4 and AvrRpm1 (Macho et al., 2010; Rufián *et al.*, 2015); and (*iii*) SAR triggered by either virulent or avirulent bacteria (Macho et al., 2010; Rufián et al., 2015). On the other hand, HopZ1a triggers ETI in Arabidopsis upon recognition by the NLR ZAR1 (HOPZ-ACTIVATED RESISTANCE 1) (Lewis *et al.*, 2010), a defense response that is independent of salicylic acid (SA) and EDS1 (Lewis et al., 2010; Macho et al., 2010).

HopZ1a belongs to the YopJ/HopZ superfamily of T3Es, which includes representatives from both animal and plant pathogens (reviewed by Ma & Ma, 2016). Many of these T3Es have been described to function as acetyltransferases, among other biochemical activities (Mittal *et al.*, 2006; Zhou *et al.*, 2005; Trosky *et al.*, 2004; Jones *et al.*, 2008; Lee *et al.*, 2015). HopZ1a has been shown to display acetyltransferase activity, with varying degrees of efficiency, on some of its interacting partners in the plant (Lee *et al.*, 2012; Jiang *et al.*, 2013; Lewis *et al.*, 2013; Zhang *et al.*, 2016; Lee *et al.*, 2019; Bastedo *et al.*, 2019). HopZ1a acetyltransferase activity is completely dependent on the integrity of the catalytic triad cysteine (C216), since a HopZ1a^C216A^ mutant behaves as a catalytically inactive mutant (Lee et al., 2012). Likewise, residue C216 is essential for all described HopZ1a virulence and avirulence functions *in planta* (Ma *et al.*, 2006; Lewis *et al.*, 2008; Macho et al., 2010; Lewis et al., 2014; Rufián et al., 2015). HopZ1a has also been described to auto-acetylate in two serine residues that are essential for HopZ1a function (Ma *et al.*, 2015) and in a lysine residue that partially contributes to HopZ1a function (Lee et al., 2012; Ma et al., 2015; Rufián et al., 2015).

A number of plant proteins have been described to interact with HopZ1a and have been proposed as targets of its virulence activity, such as isoflavone biosynthesis enzyme HID1, JASMONATE ZIM DOMAIN (JAZ) transcriptional repressors, or tubulin (Zhou *et al.*, 2011; Lee et al., 2012; Jiang et al., 2013). Additional plant proteins have been described to participate in the recognition of HopZ1a by the plant defense system, likely playing a double role as virulence targets and immunity-triggers (Bastedo et al., 2019; Liu *et al.*, 2019; Albers *et al.*, 2019). HopZ1a-dependent acetylation of the Receptor-Like Cytoplasmic Kinase (RLCK) ZED1 (HOPZ-ETI-DEFICIENT 1), a pseudokinase that acts as a decoy, is detected by the ZAR-1 resistance protein, triggering ETI (Lewis et al., 2010; Lewis et al., 2013). Resistance requires the formation of a HopZ1a-ZED1-ZAR1 complex, with ZED1 acting as an adaptor, but also involves additional RLCKs such as PBS1-like (PBL) kinases (Bastedo et al., 2019), SZE1 and SZE2 (SUPPRESSOR OF ZED) kinases (Liu et al., 2019), or the scaffold protein remorin, also associated to plant immune signaling (Albers et al., 2019). Interaction with several RLCKs has also been described for HopZ3, a homolog of HopZ1a that is present in *P. syringae* strain B728a (Lee et al., 2015).

Interestingly, the targeting of host kinases is a common theme among T3Es within the YopJ/HopZ superfamily. The *Yersinia* T3E YopJ acetylates key serine and threonine residues of several host Mitogen-Activated Protein Kinase Kinases (MAP2Ks or MKKs) and MAP kinase kinase kinases (MAP3Ks), abolishing the phosphorylation of these residues, which in turn leads to inactivation of downstream immune signaling and responses (Mittal et al., 2006; Mukherjee *et al.*, 2006; Paquette *et al.*, 2012; Meinzer *et al.*, 2012; reviewed by Ma & Ma, 2016). YopJ can also acetylate lysine residues of several of its target kinases, however this modification does not seem to be essential for its inhibitory function (Mukherjee et al., 2006; Paquette et al., 2012). Within the same superfamily, AvrA from *Salmonella* and VopA from *Vibrio* acetylate key serine, threonine, and lysine residues of their corresponding target MKKs, resulting in inhibition of kinase activity and the suppression of immune responses (Trosky *et al.*, 2007; Jones et al., 2008).

In plants, MAPK-dependent signaling networks participate in defense against pathogens, since PRR recognition of PAMPs leads to activation of MAPK modules and ultimately to the immune response (reviewed by (Pitzschke *et al.*, 2009; Feng *et al.*, 2012; Meng & Zhang, 2013). HopZ1a suppresses MAPK activation in Arabidopsis (Lewis et al., 2014), while HopZ3 directly interacts with MAPK4 (Lee et al., 2015). Further, several other *P. syringae* T3Es can suppress plant defense signaling by targeting MAPK cascades, such as HopAI1, which interferes with several MAP kinases (MPKs) (Zhang *et al.*, 2007a; Zhang *et al.*, 2012), or HopF2, which blocks phosphorylation of MKK5 (Wang *et al.*, 2010).

Although several reports have described plant targets of HopZ1a (Zhou et al., 2011; Lee et al., 2012; Jiang et al., 2013; Albers et al., 2019), the molecular mechanisms of HopZ1a-mediated suppression of immunity, particularly those regarding ETI or SAR, remain unclear. Given the broad plant defense suppression abilities of HopZ1a, the proclivity of YopJ/HopZ-family T3Es to target host kinases, the targeting of decoy pseudokinase ZED1 by HopZ1a, and the importance of MAPK cascades as general regulators of immune signaling, we considered MKKs as potential virulence targets of HopZ1a involved in its suppression of PTI, ETI and SAR. The *Arabidopsis* genome presents ten genes encoding MKKs, of which only eight are likely to be expressed (Zhang *et al.*, 2008). Among these, MKK7, MKK3, and the functionally redundant pairs MKK1/2 and MKK4/5 have all been identified as positive regulators of plant defense (Asai *et al.*, 2002; Doczi *et al.*, 2007; Zhang *et al.*, 2007b; Meng & Zhang, 2013). However, while MKK3, MKK1/2 and MKK4/5 cascades have not been associated to SAR to date, MKK7 has been shown to be essential for SAR activation (Zhang et al., 2007b). Considering the evidence available, we decided to investigate MKK7 as a potential target for HopZ1a.

In this work, we show that MKK7 contributes to PAMP-triggered callose deposition, ROS burst, and MAPK activation, contributing to resistance against *Pto* DC3000 and a non-pathogenic T3SS null mutant. We also demonstrate that MKK7 contributes to immune responses triggered by AvrRpt2. Remarkably, we found that HopZ1a interacts with MKK7, and show that bacteria-delivered HopZ1a interferes with and suppresses MKK7-dependent PTI, ETI, and SAR. Finally, we show that HopZ1a acetylates MKK7 in a conserved lysine residue, which we demonstrate to be essential for MKK7 activity, leading to a reduction of MKK7 self-phosphorylation.

## MATERIALS AND METHODS

### Bacterial strains and growth conditions

*Escherichia coli* DH5α or NCM631 (Hanahan, 1983; Govantes *et al.*, 1996), *Pseudomonas syringae* pv. tomato DC3000 (Cuppels, 1986), *Pseudomonas fluorescens* 55 (Huang *et al.*, 1988; Jamir *et al.*, 2004), *Agrobacterium tumefaciens* C58C1 (Deblaere *et al.*, 1985), and derivatives carrying a plasmid (**Table 1**), were grown at 37°C (*E. coli*) or 28°C (*Pseudomonas and Agrobacterium*) in Luria-Bertani (LB) medium (Bertani, 1951) supplemented with antibiotics when appropriate. Antibiotics were used at the following concentration: 50 μg/ml kanamycin or 50 μg/ml spectinomycin for *E. coli*, 15 μg/ml kanamycin for *P. syringae* strains, 5 μg/ml tetracycline and 15 μg/ml kanamycin for *P. fluorescens* strains, and 50 μg/ml kanamycin, 50 μg/ml rifampicin, 50 μg/ml spectinomycin, and 5 μg/ml tetracycline for *Agrobacterium*. All plates used to grow plant-extracted bacteria contained cycloheximide (2 μg/ml) to prevent fungal contamination.

**Table 1.**
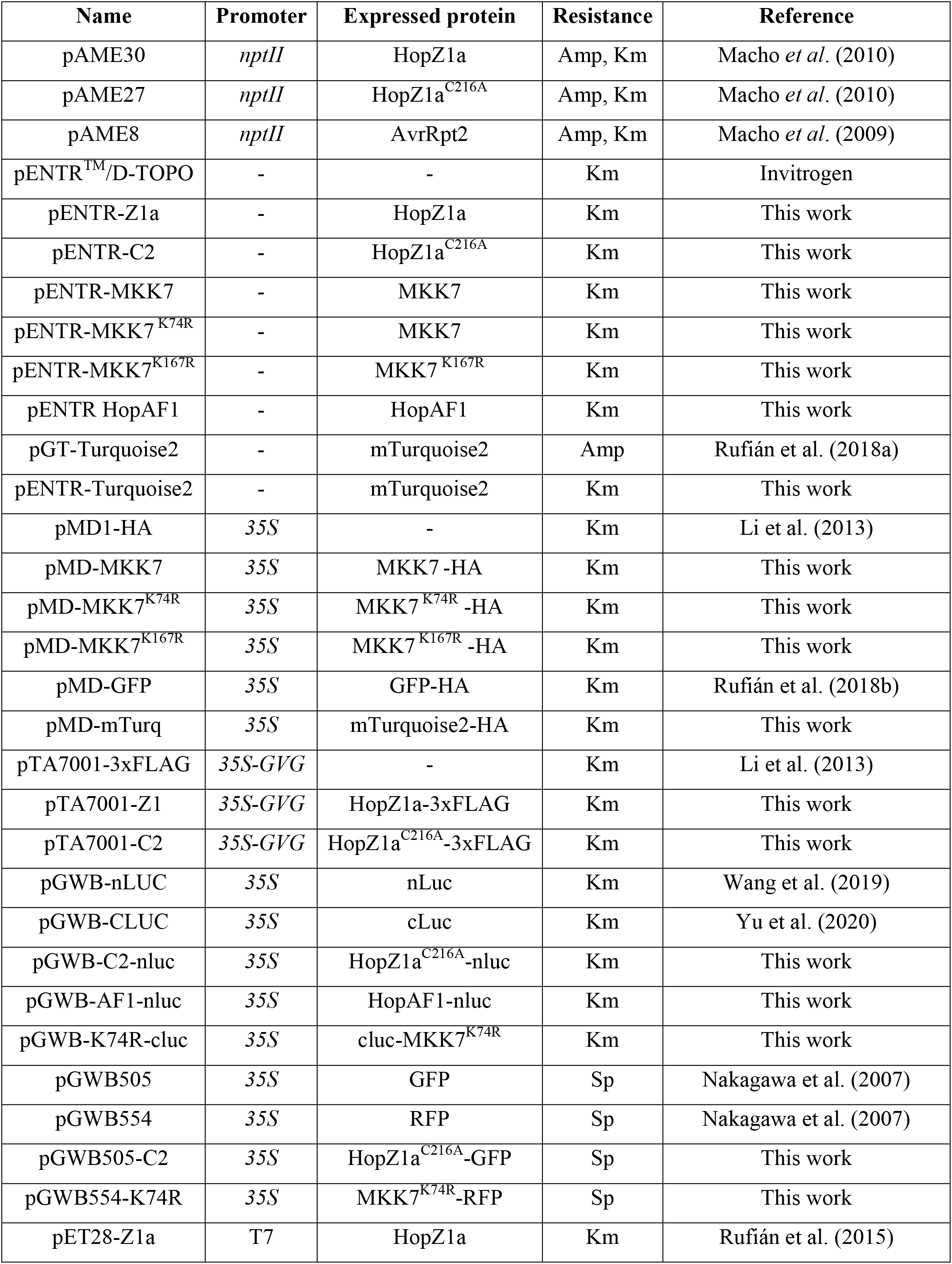
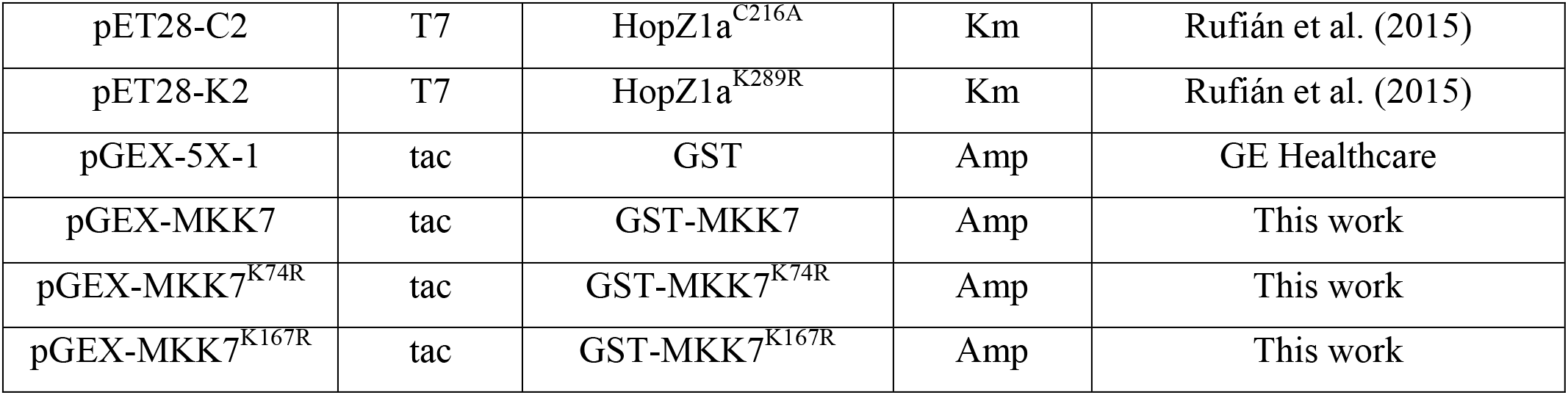
Plasmids used in this work.

### Plasmid generation

Plasmids used in this work are listed in **Table 1**. All PCRs were performed using Q5 High-Fidelity DNA Polymerase (New England Biolabs, USA), unless otherwise stated. 6xHis-HopZ1a fusion proteins were generated as previously described (Rufián et al., 2015). HopZ1a and HopZ1a^C216A^ were amplified by PCR using plasmids pAME30 and pAME27 as templates, and primers Z1pET-F and Z1pET-R (**Table 2**). PCR-amplified DNA fragments, encoding the corresponding ORFs were digested with *Nde*I and *BamH*I and cloned into the corresponding sites of expression vector pET28a(+). The resulting vectors pET28-Z1a, and pET28-C2 carry HopZ1a or HopZ1a^C216A^, 6xHis N-terminal fusion proteins, respectively.

**Table 2.**
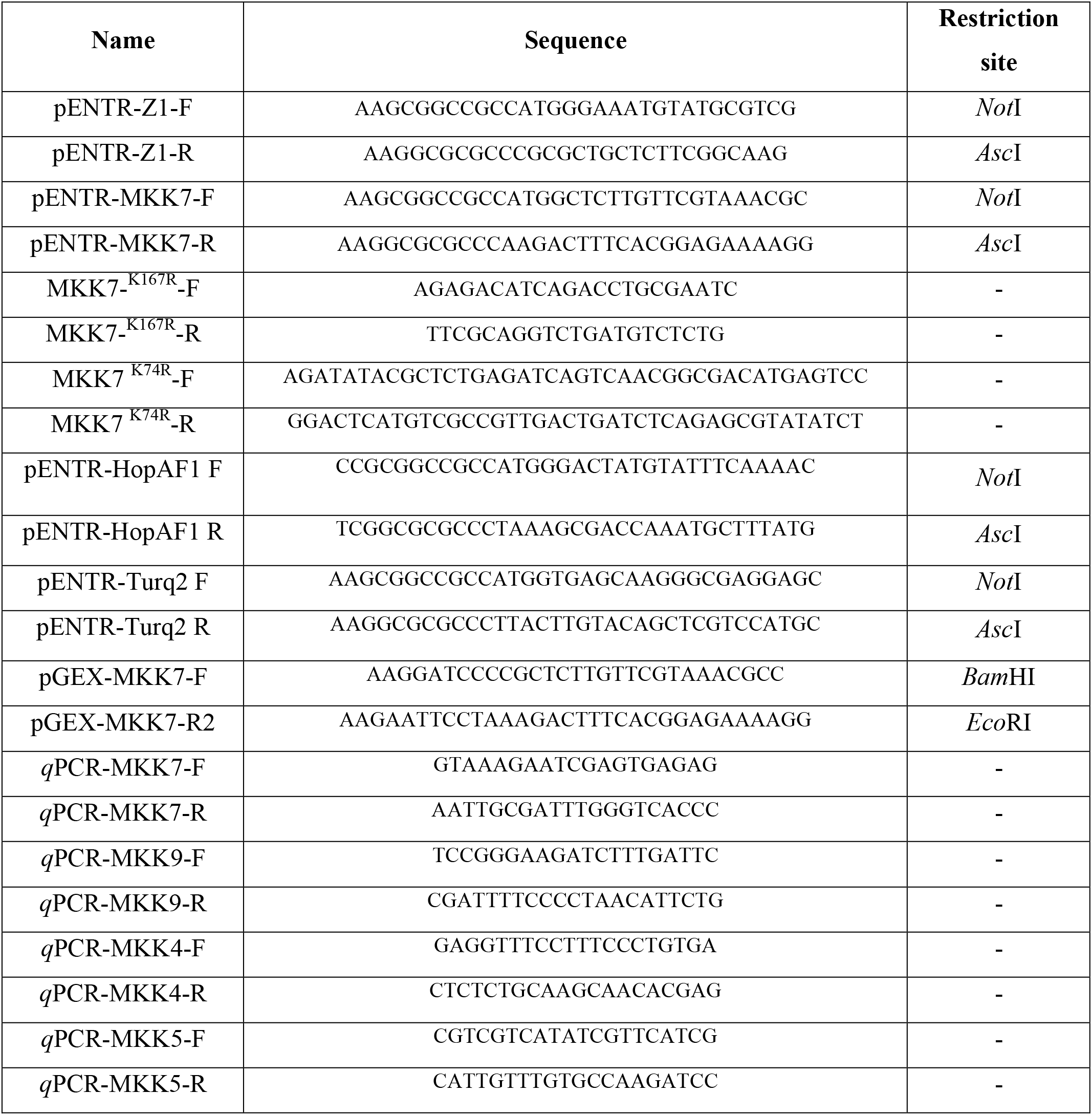
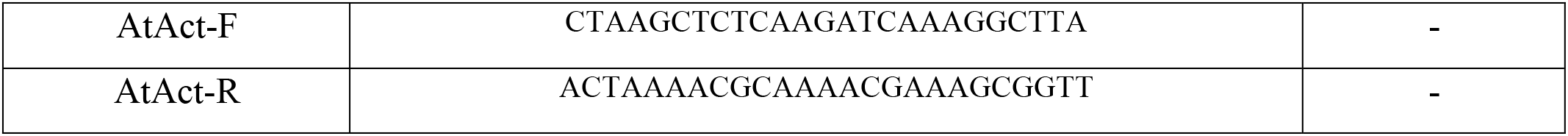
Primers used in this work.

For GST fusion proteins, MKK7 ORF was PCR amplified from Arabidopsis genomic DNA using primers MKK7-F and MKK7-R1 (**Table 2**) and cloned into pGEX-5X-1 (GE Healthcare, USA) into *Bam*HI-*Eco*RI restriction sites. MKK7 point mutants were generated following the instructions for the NZY Mutagenesis kit (NZY Tech, Portugal) using the primers listed in **Table 2**. The point mutations introduced were verified by sequencing.

To generate Gateway-cloning intermediates, HopZ1a and HopZ1a^C216A^ were amplified by PCR using plasmids pAME30 and pAME27 as templates, and primers Z1a pENTR-F and Z1a pENTR-R (**Table 2**). MKK7 was amplified from Arabidopsis genomic DNA using primers MKK7 pENTR-F and MKK7 pENTR-R (**Table 2**). All these fragments were digested with *Asc*I and *Not*I and cloned into the corresponding sites on pENTR/D (Invitrogen, USA). After validation by sequencing, we used the Gateway LR Clonase II Enzyme mix (Invitrogen, USA) to clone the fragments into their respective destination vectors (**Table 1**).

### Dexamethasone treatment

Dexamethasone (SIGMA, USA) stock was prepared at 10 mM in ethanol. To induce MKK7 expression from the dexamethasone-inducible promoter, we infiltrated leaves with either 10 uM dexamethasone solution in water (DEX) or 0.1% ethanol in water (mock). Plant discs and seedlings were incubated either with 3 ml DEX or mock solution for 24 hours.

### Plant material and bacterial inoculations

*Arabidopsis thaliana* (Col-0) and the derivatives *MKK7*-DEX, *asMKK7* (Zhang et al., 2007b) and *MKK7*-DEX/*zar1-1* were grown in soil, in temperature-controlled chambers, at 21°C with a controlled photoperiod of 8h light/16h dark with a light intensity of 200 μmol/m^2^/s. *Nicotiana benthamiana* plants were grown in soil in temperature-controlled chambers, at 21°C with a controlled photoperiod of 16h light/8h dark with a light intensity of 200 μmol/m2/s.

For *P. syringae* growth assays, bacterial lawns were grown on LB plates for 48 h at 28 °C, scrapped off the plates and resuspended into 2 mL of 10 mM MgCl_2_. The OD_600_ was adjusted to 0.1 (5 × 10^7^ cfu/mL) and serial dilutions were made to reach the final inoculum dose (5 × 10^4^ cfu/mL). MKK7-DEX plants were infiltrated with either DEX or mock solution 2 hours before bacterial inoculation. Three fully expanded leaves of 5-week-old Arabidopsis plants were inoculated using a 1-mL syringe without needle. Samples were taken from infiltrated leaves at 4 days post inoculation (dpi) using a 10-mm-diameter cork-borer (SIGMA, St. Louis, MO, USA). One disc was taken per leaf, three discs per plant, placed into 1 mL of 10 mM MgCl_2_, and homogenized by mechanical disruption. Serial dilutions of the resulting bacterial suspensions were plated onto LB plates supplemented with 2 μg/mL of cycloheximide.

Competitive index assays were performed as previously described for Arabidopsis (Macho *et al.*, 2016). Briefly, fully expanded leaves of 4-5 weeks-old plants were infiltrated with a 1:1 suspension of the indicated bacteria at 5×10^4^ cfu/ml. Four days later, bacteria were recovered as described above. Serial dilutions were spread on LB plates, and colonies were replica-plated onto plates containing the appropriate antibiotic and without antibiotic. Index value was calculated as the strain carrying the effector-to-wild type ratio within the output sample divided by the strain carrying the effector-to-wild type ratio within the input (inoculum).

For measuring SAR, we used a protocol previously described in Rufian et al. (2019) with modifications. Briefly, three fully expanded leaves of five weeks-old plants were infiltrated either with DEX or mock solution. Two hours after treatment, same leaves were infiltrated either with 10 mM MgCl_2_ (mock), or a 5×10^5^ cfu/ml bacterial suspension of either *P. fluorescens* pLN18 or *P. fluorescens* expressing HopZ1a. Three days after inoculation, three distal leaves were inoculated with a 5×10^4^ cfu/ml suspension of DC3000 prepared as described above. Four days after DC3000 inoculation, tissue was collected as described above.

Results presented in all growth assays are the mean of at least three replicates showing typical results from at least three independent experiments. Errors bars represent standard error. Each value was analyzed using a homoscedastic and 2-tailed Student’s t-test and the null hypothesis: mean index is not significantly different from 1. Pair-wise comparisons were similarly analyzed with the null hypothesis of mean indexes not significantly different (P value < 0.05). For comparing more than two mean values statistical significance was tested using one-way ANOVA (α=0.05) with Tukey’s multiple comparisons test.

Transient expression assays in *N. benthamiana* were performed as previously described (Rufián et al., 2015). Briefly, 5-week old plants were infiltrated with an *Agrobacterium* C58C1 solution at OD_600_ 0.5 in 10 mM MgCl_2_, 10 mM MES (SIGMA, USA), 200 μM 3′,5′-Dimethoxy-4′-hydroxyacetophenone (acetosyringone) (SIGMA, USA) carrying the corresponding binary plasmids (**Table 1**). Strains carrying the indicated plasmids were co-inoculated at an OD600 of 0.5 each. When required, plants were treated with DEX 24 h after agroinfiltration. Samples were collected (for immunoprecipitation assays) or photographed (for development of macroscopic HR, luciferase assays, and subcellular localization) at 48 hpi.

### RNA extraction and RT-*q*PCR

For RNA extraction assays, we used 100 mg of the corresponding plant tissue, seedlings or adult leaves, previously treated with either DEX or mock solution, frozen and grounded into liquid nitrogen. RNA was extracted using TRISURE (Bioline, UK) according to instructions from the manufacturer. Reverse transcriptase (RT) reactions were performed using iScript cDNA Synthesis Kit (Bio-Rad, USA) and 1 μg of total RNA, previously treated with DNaseI (Takara, Japan).

For RT-*q*PCRs, 2 μl (for *MKK7* quantification) or 1 μl (for the rest) of a 1/2 dilution of the cDNA was added to a reaction containing 1X of SsoFast EvaGreen (Bio-Rad, USA), 0.5 μM Forward and Reverse primers (**Table 2**). Reactions were performed in a CFX96 thermocycler (Bio-Rad, USA) with a first denaturing step at 95°C 1 min, and 45 cycles of 95°C 10 s and 60°C 15 s. In all cases, At*ACT2* was used as housekeeping gene. Relative expression was calculated using the 2^−ΔΔCt^ method (Livak & Schmittgen, 2001).

### Measurement of ROS generation

Oxidative burst was quantified as previously described (Sang & Macho, 2017). Plant discs were incubated overnight with either DEX or mock solution. ROS was elicited with 100 nM flg22 (GeneScript, USA). Twenty leaf discs from 4 weeks-old plants were used for each condition. Luminescence was measured using a GloMax 96 Microplate Luminometer (Promega, USA).

### Callose deposition

Callose deposition was detected by aniline blue staining as described by Adam and Somerville (1996). Leaves from 4 weeks-old plants were infiltrated either with DEX or mock solution. Twenty-four h after treatment, the same leaves were infiltrated with 100 nM flg22 (for flg22-induced callose deposition experiments) or with the indicated bacterial strain at 5×10^7^ cfu/ml (for HopZ1a-interference assays). Samples were collected 18 hours after the second inoculation and stained with aniline blue. Three plants were used for each treatment, using 2 leaves per plant, and taking two representative images per leaf. Callose deposition was measured using Fiji distribution of ImageJ.

### Plant protein extraction

For MAP Kinase activation assays, 10 days-old seedlings incubated in liquid MS were treated with either DEX or mock solution. Twenty-four hours later, treatment solutions were removed, and seedlings were immersed in a 100 nM flg22 solution. Ten seedlings were collected for each sample and treatment at the indicated time points, frozen in liquid nitrogen and homogenized using a Tissue Lyser (QIAGEN, Germany). Proteins were extracted in 100 μl extraction buffer (100 mM TRIS pH 7.5, 150cmM NaCl, 5 mM EDTA, 10% glycerol, 1x protease inhibitor cocktail (SIGMA, USA) and 1x phosphatase inhibitor (Cell Signaling, USA) and centrifuged 10 minutes at 16,000 *g*. The resulting protein samples were quantified using a BioRad protein assay (BioRad, USA). For SDS-PAGE,10 μg of protein was loaded per sample.

For PR1 accumulation assays in local tissue, two fully expanded Arabidopsis young leaves were treated either with DEX or mock solution. Two hours later, the same leaves were inoculated with either 10 mM MgCl_2_ (mock) or a 5×10^5^ cfu/ml bacterial solution. Samples were taken 2 days after inoculation and proteins were extracted as described above. For PR1 detection in systemic tissue, three fully expanded leaves of 5-week-old plants were infiltrated with either DEX or mock solution. Two hours after treatment, the same leaves were infiltrated with either 10 mM MgCl_2_ solution (mock), or a 5×10^5^ cfu/ml bacterial suspension of either *P. fluorescens* pLN18, or *P. fluorescens* expressing HopZ1a. Three days after infiltration, samples were taken from systemic leaves, and proteins were extracted as described.

For immunoprecipitation assays, samples were taken from inoculated leaves at 2 dpi using a 7.5-mm-diameter cork-borer (SIGMA, USA). Forty leaf discs were taken per sample and homogenized into liquid nitrogen. Proteins were extracted using 1.6 ml of extraction/washing buffer (50 mM Tris-HCl pH 7.5, 150 mM NaCl, 0.1% Triton, 0.2% NP-40, 6 mM BME and 1x protease inhibitor cocktail [SIGMA, USA]) during 30 minutes at 4°C. The resulting homogenate was centrifuged at 16,000 *g* for 30 min at 4°C, and the supernatant was filtered using Micro Bio-spin™ Chromatography Columns (Bio-Rad, USA) and collected into a new tube. Twenty μl of pre-equilibrated anti-HA agarose (SIGMA, USA) were added to each tube and incubated for 3 hours at 4°C in an end-over-end shaker. After incubation, tubes were spin down (900 *g*) and beads washed 3 times. To elute the proteins, beads were boiled 5 min at 95°C in 20 μl 3x Laemmli buffer.

### Western hybridizations

Samples were resolved on 10% acrylamide SDS-PAGE gels using a Mini protean system (Bio-Rad, USA) and transferred to PVDF Immobilon-P membranes (Millipore, USA), using the Semi-Dry Transfer System (Bio-Rad, USA) during 30 minutes at 15 V. Western blots were carried out using standard methods, and 1:5000 dilution of primary anti-His (SIGMA, USA), 1:10000 for anti-GST (SIGMA, USA), 1:5000 for anti-HA (SIGMA, USA), 1:5000 for anti-FLAG (SIGMA, USA), 1:5000 for anti-Luciferase (SIGMA, USA), 1:1000 for anti-Tubulin (Abiocode, USA), 1:5000 for anti-MPKs (Cell Signaling Biotechnology, USA) or 1:5000 for anti-PR1 ((Wang *et al.*, 2005), a kind gift from P. Tornero, IBMCP-CSIC). For secondary antibodies, we used 1:10000 dilution of a secondary Anti-Rabbit antibody (SIGMA, USA) or 1:80000 for Anti-Mouse antibody (SIGMA, USA). Membranes were developed using the Bio-Rad Clarity Western ECL Substrate (Bio-Rad, USA) following instructions from the manufacturer.

### Luciferase assays

Split-LUC assays were performed as previously described (Yu *et al.*, 2020). Generally, *A. tumefaciens* strains containing the indicated plasmids (**Table 1**) were infiltrated into *N. benthamiana* leaves. Two days after inoculation, leaves were infiltrated with 0.5 mM luciferin in water, and kept in the dark for 5 min before CCD imaging. Images were taken with VersArray 1300B (Roper Scientific, USA). Protein accumulation was determined by immunoblot as described above.

### Förster Resonance Energy Transfer-Fluorescence Lifetime Imaging (FRET-FLIM)

FRET-FLIM experiments were performed as previously described (Rosas-Diaz *et al.*, 2018), with several modifications. Briefly, AtMKK7^K74R^ (fused to RFP, acceptor) was expressed from plasmid pGWB554, and either HopZ1a or HopZ1a^C216A^ (fused to GFP, donor) from plasmid pGWB505. FRET-FLIM experiments were performed on a Leica TCS SMD FLCS confocal microscope excitation with WLL (white light laser) and emission collected by a SMD SPAD (single photon-sensitive avalanche photodiodes) detector. Twenty-four hours after infiltration, *N. benthamiana* plants transiently co-expressing donor and acceptor proteins were visualized under the microscope. Accumulation of the GFP- and RFP-tagged proteins was estimated before measuring lifetime. Tuneable WLL set at 488 nm with a pulsed frequency of 40 MHz was used for excitation, and emission was detected using SMD GFP/RFP Filter Cube (with GFP: 500-550 nm). The fluorescence lifetime shown in the figures, corresponding to the average fluorescence lifetime of the donor, was collected and analyzed by PicoQuant SymphoTime software. Mean lifetimes are presented as mean ± SEM based on eight images from three independent experiments.

### Conductivity Assays

To measure cell death induced by *P. syringae* strains during an incompatible interaction associated to the onset of HR in Arabidopsis, Col-0, and asMKK7 leaves were syringe-infiltrated with a 5 × 10^7^ cfu/ml suspension of the indicated strain, and four discs were taken per leaf at the indicated time points and immersed in 6 ml of distilled water for 30 min. To measure cell death induced by *Agrobacterium*-mediated MKK7, MKK7^K74R^, or MKK7K167R expression in *N. benthamiana*, leaves were syringe-infiltrated as indicated above. Two hours after infiltration, four leaf discs were immersed in 6 ml of distilled water for 30 min. In all assays, leaf discs were then transferred to 6 ml of distilled water and conductivity was measured at the indicated time point using a portable conductivity meter Crison CM35 (Hach-Lange, Spain).

### Protein expression and purification *in vitro*

Plasmids used in this work are listed in **Table 1**. All proteins were expressed in *E. coli* NCM631 after induction with 0.1 mM IPTG at 20°C. His-tagged proteins were purified using Ni-NTA agarose (Quiagen, USA). GST-tagged proteins were purified using Glutathione Sepharose 4B (GE Healthcare, USA). Protein concentration was determined using Bio-Rad protein assay (Bio-Rad, USA).

### *In vitro* acetylation assay

For ^14^C-based acetylation assays, 3 μg of the effector (6xHis-HopZ1a, 6xHis-HopZ1a^C216A^ or 6xHis-HopZ1a^K289R^) and 5 μg of the substrate (GST-MKK7 or GST-MKK7^K167R^) were incubated in acetylation buffer containing 50 mM HEPES pH 8, 10% Glycerol, 10 mM Sodium butyrate, 1 mM DTT and 1 mM PMSF, with 100 nM inositol hexakisphosphate (IP6 or phytic acid, SIGMA, USA) and 22 nCi Acetyl-CoA (Perkin Elmer, USA). The reaction was incubated for 1 hour at 30°C, then stopped by adding Laemmli buffer and boiled at 95°C for 5 minutes. Twenty μl of each sample were separated by SDS-PAGE, and proteins were transferred onto a PVDF membrane. Acetylation was detected by autoradiography. As a loading control 20 μl of the same samples were separated by SDS-PAGE and stained with Coomassie blue.

### *In vitro* kinase assay

For *in vitro* kinase assays, 1 μg of GST-MKK7, GST-MKK7^K74R^, or GST-MKK7^K167R^ were incubated into phosphorylation buffer containing 50 mM Tris-HCl pH 7.4, 5 mM MnCl_2_, 5 mM MgCl_2_, 1 mM DTT, 1 μM cold ATP and 5 μCi [γ^32^P] - ATP (Perkin Elmer, USA). The reaction was incubated for 30 minutes at 30°C and stopped by adding Laemmli buffer and boiling at 95°C for 5 minutes. Ten μl of each sample were separated by SDS-PAGE, and phosphorylation detected by autoradiography. As a loading control, 10 μl of the same samples were separated by SDS-PAGE and stained with Coomassie blue.

## RESULTS

### HopZ1a interacts with MKK7 *in planta*

Considering the published evidence, we decided to investigate MKK7 as a potential target for HopZ1a, using *Agrobacterium*-mediated transient expression in *Nicotiana benthamiana*. Since expression of MKK7 has been shown to trigger cell death in *N. benthamiana* (Popescu *et al.*, 2009), we used a catalytically inactive version of this protein (MKK7^K74R^) that lacks kinase activity due to a defect on ATP binding (Dai *et al.*, 2006; Zhang et al., 2007b). As for HopZ1a, we used both the wild type version and a catalytically inactive version (HopZ1a^C216A^) that lacks acetyltransferase activity and does not trigger cell death when transiently expressed *in planta* (Ma et al., 2006; Lewis et al., 2008; Jiang et al., 2013; Lewis et al., 2013).

We first co-expressed MKK7^K74R^-HA (or GFP-HA, as a control) with either HopZ1a-3xFLAG or HopZ1a^C216A^-3xFLAG in *N. benthamiana* leaves, and performed co-immunoprecipitation (coIP) using anti-HA beads. The results show that both HopZ1a-3xFLAG and HopZ1a^C216A^-3xFLAG associate with MKK7^K74R^-HA, but not with the GFP-HA control (**Fig. 1a**). Expression of wild type HopZ1a, but not HopZ1a^C216A^, eventually triggered cell death in *N. benthamiana*, at variance with a recent report (Baudin *et al.*, 2017), a difference likely due to higher expression levels being achieved in our experimental settings. Thus, we decided to further confirm the interaction detected by coIP using only HopZ1a^C216A^, to benefit from a wider time range for sampling. To this end, we performed split-luciferase complementation assays (**Fig. 1b**), in which MKK7^K74R^ and HopZ1a^C216A^ were fused to the C-terminal (cluc) or N-terminal (nluc) domains of the luciferase protein, respectively (**Table 1**). As a negative control, we used the *P. syringae* effector HopAF1, which, as HopZ1a, is associated to the plasma membrane through acylation (Lewis et al., 2008; Washington *et al.*, 2016). The results show a strong luminescence signal in tissues expressing MKK7^K74R^-cluc and HopZ1a^C216A^-nluc, and only background signal for MKK7^K74R^-cluc and HopAF1-nluc (**Fig. 1b**), confirming the interaction between MKK7 and HopZ1a. To further analyze the direct interaction between MKK7^K74R^ and HopZ1a *in planta*, we used Förster Resonance Energy Transfer-Fluorescence Lifetime Imaging (FRET-FLIM), using HopZ1a^C216A^ fused to GFP, and MKK7^K74R^ fused to RFP. The results show that the GFP fluorescence lifetime for the HopZ1a^C216A^-MKK7^K74R^ co-expression was significantly shorter than that of the control samples, as expected from a direct interaction between MKK7 and HopZ1a (**Fig. 1c**).

**FIGURE 1.**
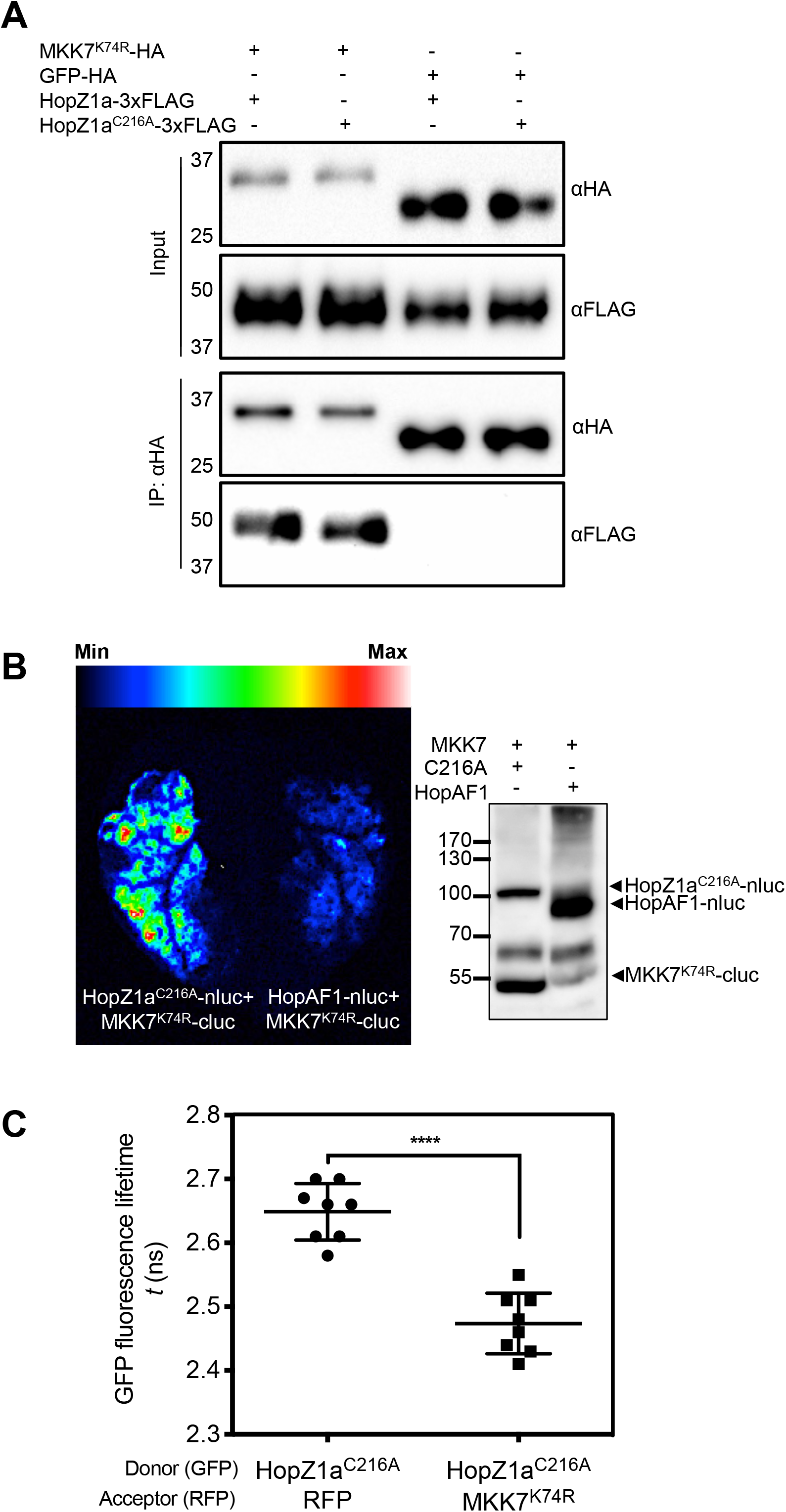
HopZ1a interacts with MKK7 *in planta.* (**A**) Co-immunoprecipitation assays using anti-HA beads. *N. benthamiana* leaves were co-inoculated with a 1:1 mix of *A. tumefaciens* carrying the indicated constructs. Forty-eight hours after infiltration, proteins were extracted and incubated with anti-HA conjugated beads. The immunoprecipitated proteins were resolved by SDS-PAGE and analyzed by Western blot with the indicated antibodies. The experiment was repeated three times with similar results. (**B**) Split-luciferase complementation assay in *N. benthamiana.* Leaves were co-inoculated with a 1:1 mixture of *A. tumefaciens* containing MKK7^K74R^-cluc with either HopZ1a^C216A^-nluc or HopAF1-nluc as a negative control. Luminescence was quantified 48 hpi and protein accumulation was determined by immunoblotting using anti-luciferase antibody. The experiment was repeated three times with similar results. (**C**) FRET-FLIM assay using *N. benthamiana* leaves inoculated with a 1:1 mixture of *A. tumefaciens* containing the indicated constructs. HopZ1a^C216A^-GFP was used as donor protein and MKK7^K74R^-RFP or free RFP (negative control) as acceptor protein. Images were taken 24-30 hpi. Lines represent average values (n=8) and error bars represent standard error. Asterisks indicate significant differences as established by Student’s test (p < 0.0001). Individual values are also shown. The experiment was repeated three times with similar results.

#### AtMKK7 participates in flg22-triggered ROS burst, MAPK activation, and callose deposition

AtMKK7 activity has been linked to basal resistance (Zhang et al., 2007b). Since HopZ1a has been shown to block the production of Reactive Oxygen Species (ROS) upon recognition of bacterial PAMP flagellin (Lewis et al., 2014), we first investigated whether *MKK7* is expressed in Arabidopsis in response to the flagellin elicitor peptide flg22 (**Fig. S1**). Results indicate that flg22 induces endogenous expression of *MKK7* (**Fig. S1**). Then, we used transgenic Arabidopsis plants expressing the *AtMKK7* gene under the control of the dexamethasone (DEX)-inducible promoter (hereafter *MKK7*-DEX plants; (Zhang et al., 2007b) to assay ROS production in response to flg22. Flg22-triggered ROS burst was significantly stronger in plants overexpressing *MKK7*, but not MKK7^K74R^ (**Fig. 2a** and **S2a**), supporting the notion that MKK7 contributes to flg22-triggered ROS production in a kinase activity-dependent manner. We also analyzed ROS production in plants expressing an antisense *MKK7* transgene under the control of a 35S promoter (hereafter, asMKK7 plants), in which *MKK7* expression is specifically compromised (Dai et al., 2006)**; Fig. S1**), ROS production after flg22 treatment on asMKK7 transgenic plants was significantly weaker than that observed in flg22-treated Col-0 plants (**Fig. 2b** and **S2b**). Altogether, these results indicate that MKK7 is involved in flg22-triggered signaling leading to the production of ROS.

**FIGURE 2.**
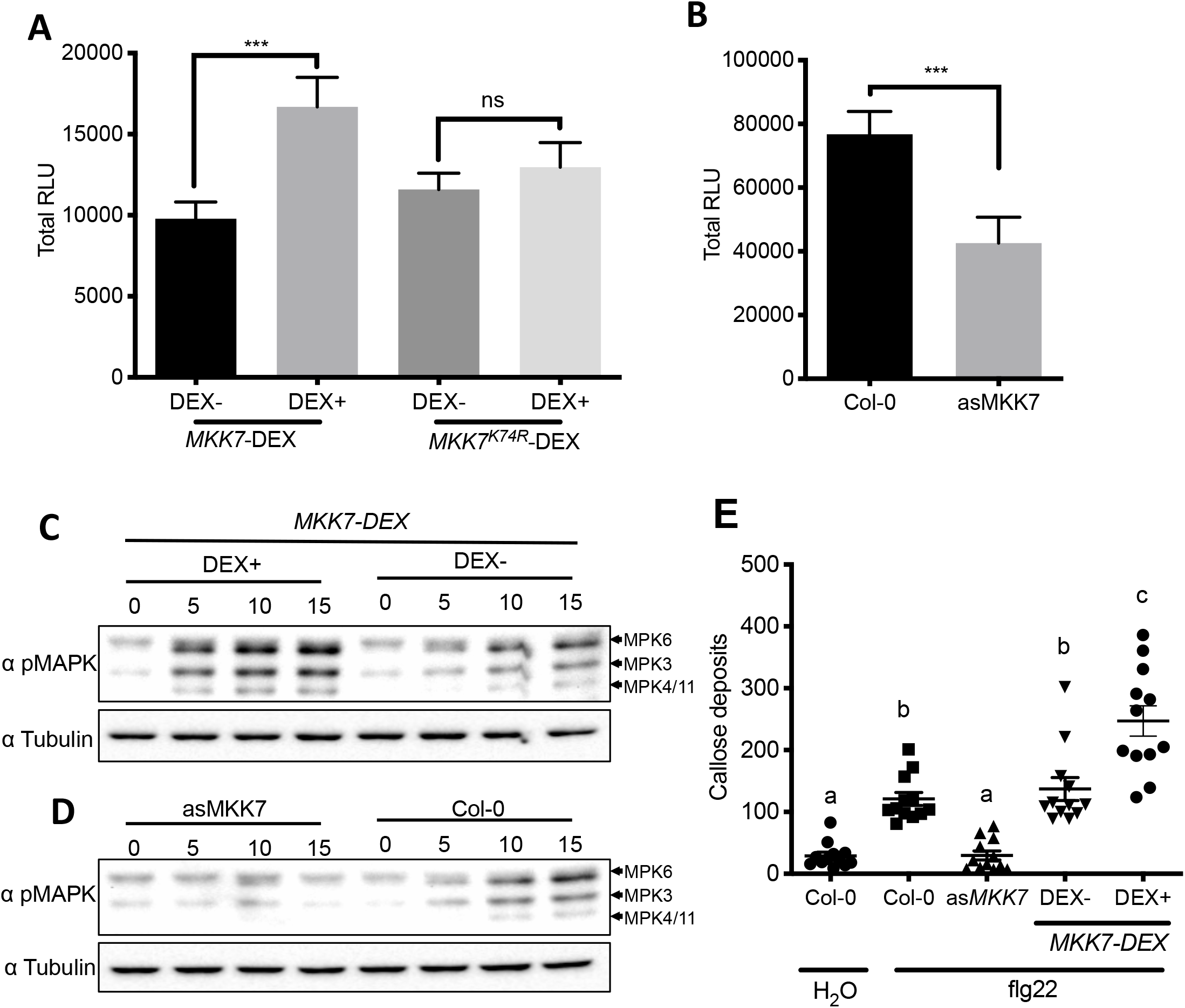
*AtMKK7* participates in flg22-triggered defence response. (**A, B**) ROS production after treatment with 100 nM flg22 of MKK7-DEX and MKK7^K74R^-DEX plants (**A**) or Col-0 and asMKK7 plants (**B**), measured in a luminol-based assay and represented as relative luminescence units (RLU). MKK7-DEX and MKK7^K74R^-DEX plants were treated either with a DEX or mock solution 24 hours before flg22 treatment. Graphs show accumulated RLU production during 40 minutes after 100 nM flg22 treatment. Error bars represent standard error (n=20). Asterisks indicate significant differences compared to the control as established by a Student’s t test (p ≤0.001). (**C, D**) MAP Kinase activation assay after 100 nM flg22 treatment. Samples were taken at four different time points (0, 5, 10 and 15 minutes, as indicated) after treatment with 100 nM flg22. MAPK phosphorylation was detected by Western blot analysis. Anti-tubulin was used as loading control. In (**C**), plants were pre-treated with either a DEX or mock solution 24 hours before flg22 treatment. (**E**) Quantification of callose deposits after flg22 treatment. MKK7-DEX plants were infiltrated with either DEX (DEX+) or mock solution (DEX−) as indicated in the figure. Twenty-four hours later, MKK7-DEX plants, asMKK7 plants, and Col-0 control plants were infiltrated with 100 nM flg22. Col-0 plants were also infiltrated with water (negative control). Eighteen hours after treatment, leaves were stained with aniline blue and observed under UV fluorescence microscope. Lines represent average values (n=12) and error bars represent standard error. Individual values are also shown. Statistical differences were determined using one-way ANOVA (α= 0.05) with Tukey’s multiple comparisons test and different letters indicate statistical significance. The experiment was repeated three times with similar results.

PRR recognition of bacterial PAMPs also leads to the activation of MAPK cascades (Boller & Felix, 2009). Transgenic expression of HopZ1a in *Arabidopsis* has been described to suppress flg22-triggered phosphorylation of MPK3 and MPK6 (Lewis et al., 2014). Interestingly, the flg22-triggered activation of MPK3, MPK6 and MPK4/11 was accelerated and enhanced in plants overexpressing MKK7 (**Fig 2c**), while it was abolished in plants expressing *asMKK7* (**Fig 2d**), indicating that MKK7 also contributes to flg22-triggered MAPK activation.

As a late PTI response, we also monitored flg22-triggered callose deposition. In agreement with the results obtained for other PTI responses, flg22-triggered deposition of callose was enhanced upon *MKK7* overexpression (**Fig. 2e**). Accumulation of callose in flg22-treated asMKK7 plants was significantly lower than flg22-treated, but not H_2_0-treated, control plants (**Fig. 2e**). Our results show that MKK7 contributes to the onset of PTI.

#### MKK7 contributes to the basal defense against P. syringae pv. tomato

Plant recognition of bacterial PAMPs ultimately leads to the restriction of bacterial growth (Zipfel *et al.*, 2004). The role played by MKK7 in plant defense against bacterial pathogens has been previously shown by monitoring the effect of MKK7 overexpression, or MKK7 silencing in Arabidopsis plants inoculated with *P. syringae* patovar maculicola or *Xanthomonas campestris* (Zhang et al., 2007b). We found that Arabidopsis plants overexpressing MKK7 also displayed enhanced resistance to *Pto* DC3000 (**Fig. 3a**), whereas asMKK7 plants reciprocally showed enhanced susceptibility compared to their respective control plants (**Fig. 3b**). Additionally, we analyzed the growth of a *Pto* DC3000 Δ*hrcV* mutant strain, which lacks a functional T3SS, and is thus non-pathogenic in *Arabidopsis* because it cannot suppress activation of PTI. Strikingly, growth of *Pto* Δ*hrcV* mutant bacteria in MKK7-overexpressing plants was less than half of that achieved in control plants (**Fig. 3c**). Taken together, these results demonstrate that MKK7 is involved in the activation of basal defenses that limit *Pto* DC3000 colonization of Arabidopsis.

**FIGURE 3.**
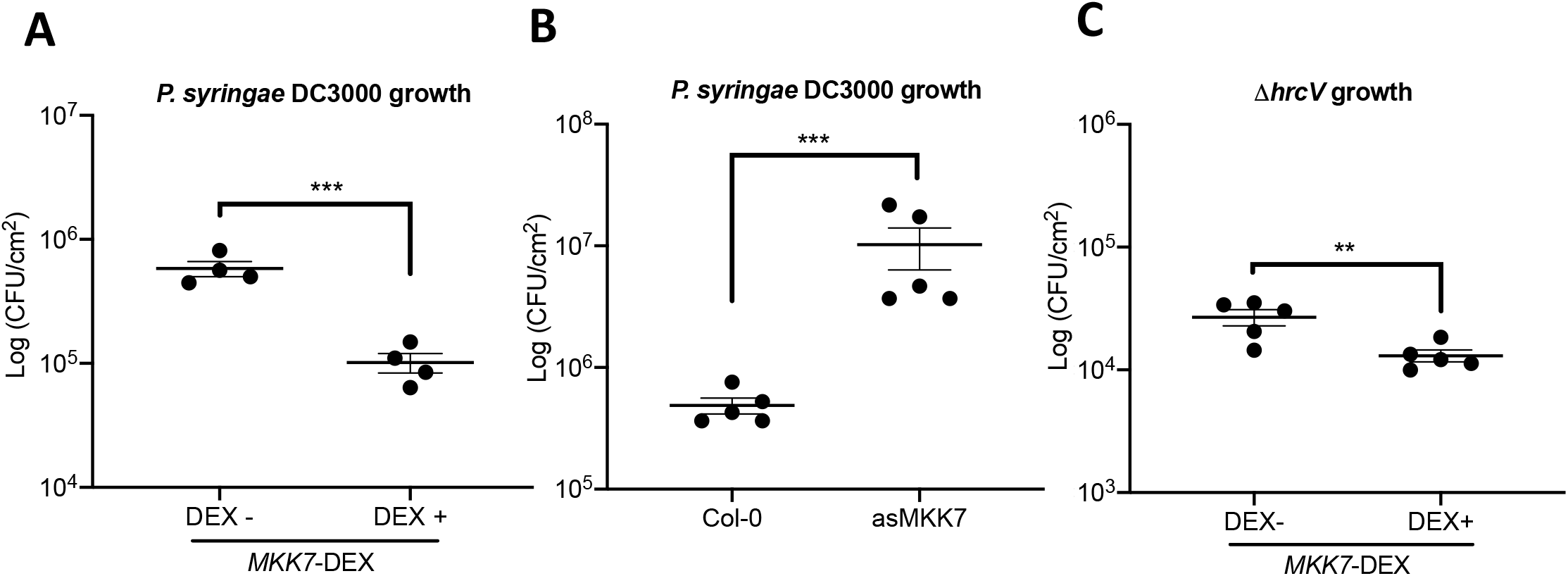
*AtMKK7* expression restrict bacterial growth. Bacterial multiplication assay in MKK7-DEX (**A, C**) or Col-0 and asMKK7 (**B**) plants. Leaves were syringe-infiltrated with a 5 × 10^4^ cfu/ml suspension of *Pto* DC3000 (**A, C**) or *Pto* Δ*hrcV* (**B**), as indicated in each graph. MKK7-DEX plants were treated with either DEX (DEX+) or Mock solution (DEX−) 2 hours before bacterial inoculation. The graphs show bacterial titers at 4 dpi. Lines represent mean values and error bars represent standard error (n=5). Individual values are also shown. Asterisks indicate significant differences as established by Student’s test (two asterisks: p < 0.01, three asterisks: p < 0.001)

#### MKK7 participates in AvrRpt2-triggered disease resistance in Arabidopsis

Expression of MKK7 is induced in Arabidopsis by inoculation of *Pto* DC3000 expressing the heterologous effector AvrRpt2 (Zhang et al., 2007b). Modification of RIN4 by AvrRpt2 triggers NLR RPS2-mediated ETI that restricts bacterial growth (Axtell & Staskawicz, 2003; Mackey *et al.*, 2003). To investigate whether MKK7 is involved in the signaling cascade of AvrRpt2-triggered immunity in Arabidopsis, we measured the effect of MKK7 silencing on attenuation of bacterial growth of DC3000 expressing AvrRpt2. To this purpose, we performed competitive index (CI) assays (Macho et al., 2016) by co-inoculating *Pto* DC3000 and *Pto* DC3000 expressing AvrRpt2, in both Col-0 and asMKK7 Arabidopsis plants (**Fig. 4a**). The CI was calculated as the *Pto* DC3000AvrRpt2/ *Pto* DC3000 ratio in the output sample, divided by this ratio within the inoculum (input), which should be close to one. As previously described for Col-0 wild type plants (Macho *et al.*, 2007), the CI of *Pto* DC3000 expressing AvrRpt2 was significantly smaller than one, showing a clear growth attenuation associated to AvrRpt2 expression. When a similar assay was carried out in asMKK7 plants, the CI obtained was 10-fold higher than the one obtained in Col-0 wild type plants and close to one, indicating that *Pto* DC3000 bacteria expressing AvrRpt2 multiplied in this plants to levels similar to those of the co-inoculated *Pto* DC3000 bacteria (**Fig. 4a**). In contrast, growth attenuation of *Pto* DC3000 expressing HopZ1a in Col-0 wild type plants, as a result of ZAR1-mediated immunity, was not significantly altered within asMKK7 plants (**Fig. 4a**). Taken together, these results indicate that MKK7 has a strong contribution to RPS2-mediated immunity, which is SA-dependent, but is not involved in SA-independent ZAR1-mediated immunity.

**FIGURE 4.**
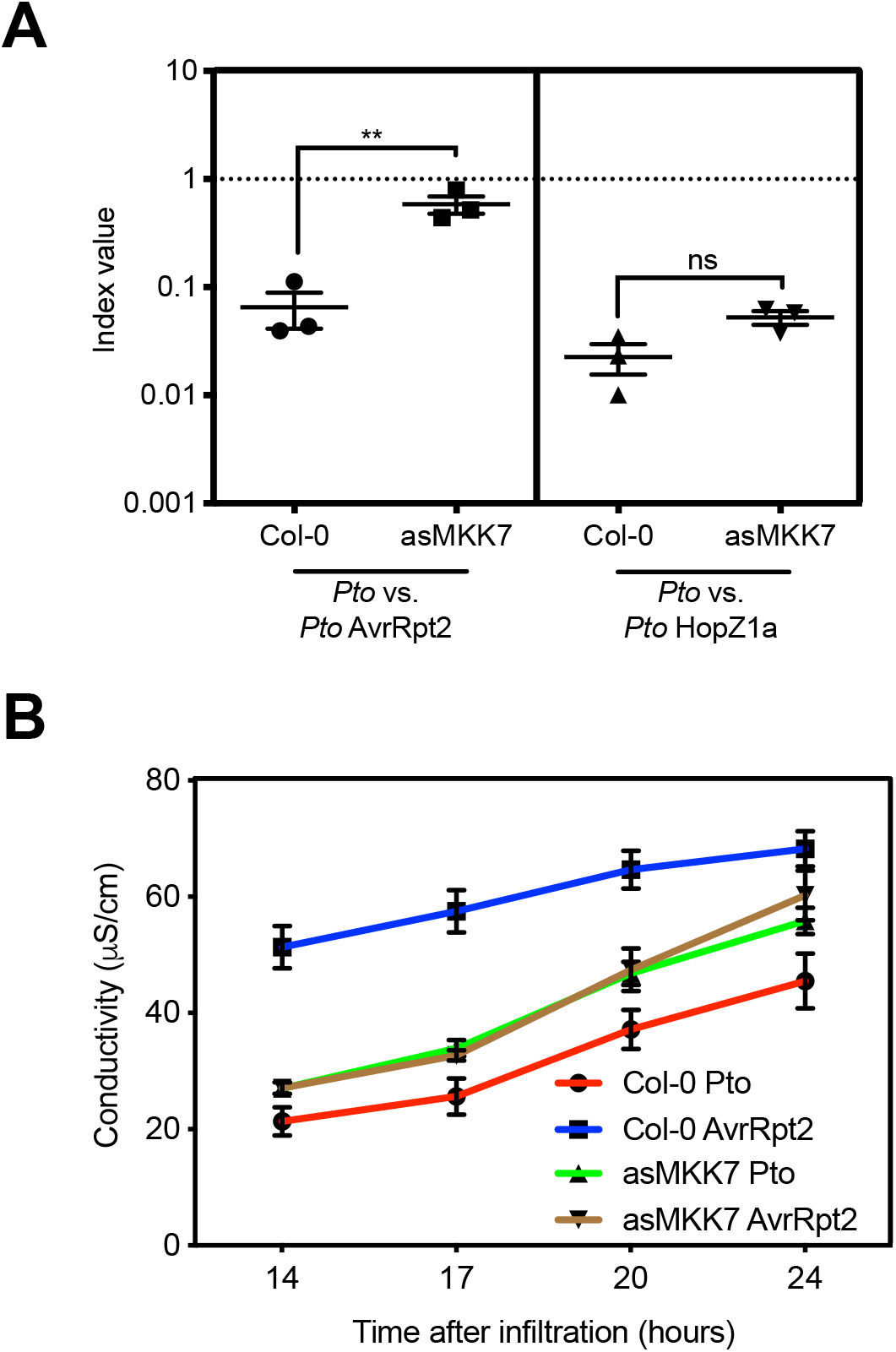
*AtMKK7* participates in AvrRpt2-mediated ETI defence response. (**A**) Competitive index (CIs) measuring bacterial proliferation in Col-0 or asMKK7 plants. *Arabidopsis* leaves were inoculated with a 1:1 mix of either *Pto* expressing AvrRpt2 and *Pto* DC3000 or *Pto* expressing HopZ1a and *Pto* DC3000 at 5 × 10^4^ cfu/ml. Bacterial loads were determined 4 dpi. CIs are calculated as the output ratio between the strain expressing the effector and the corresponding wild type or mutant strain, divided by their input ratio. Each CI mean represents three biological replicates per treatment. Individual values are shown. Error bars represent the standard error, and asterisks indicate significant differences as established by Student’s *t* test (p<0.01). Results presented are representative of three independent experiments. (**B**) Ion leakage assays in Col-0 and asMKK7 *Arabidopsis* leaves inoculated with a 5 × 10^7^ cfu/ml suspension of *Pto* DC3000 or *Pto* expressing AvrRpt2. Conductivity was measured at the indicated time points. Lines represent mean values. Error bars represent standard error (n=3).

The results obtained quantifying bacterial multiplication within mixed infections were further supported by measuring ion leakage from plant tissues, which is indicative of the activation of HR-associated cell death. To this purpose, we performed individual bacterial inoculations with *Pto* DC3000 or *Pto* DC3000 expressing AvrRpt2, in both wild type and asMKK7 Arabidopsis plants, and quantified conductivity to determine ion leakage in samples taken along a 24-hour time-course, as an indicator of the hypersensitive response (HR) triggered against AvrRpt2 (**Fig. 4b**). In the Col-0 wild type background, samples inoculated with *Pto* DC3000 expressing AvrRpt2 displayed the highest conductivity measurements at all time points, consistent with the induction of HR, while samples inoculated with *Pto* DC3000 displayed the lowest, consistent with a compatible interaction, as expected (**Fig. 4b**). Interestingly, in the asMKK7 background, samples inoculated with *Pto* DC3000 displayed conductivity levels indistinguishable from those inoculated with *Pto* DC3000 expressing AvrRpt2, and only slightly higher than those inoculated with DC3000 in the Col-0 background (**Fig. 4b**). Thus, ion leakage measurements confirm the results obtained using CIs, supporting that MKK7 has a strong contribution on RPS2-mediated immunity.

#### HopZ1a suppresses MKK7-dependent basal defence signalling in Arabidopsis

To analyze whether HopZ1a activity suppresses MKK7-dependent defense signaling in Arabidopsis, we first investigated whether HopZ1a expression from *Pto* DC3000 was able to rescue the growth attenuation caused by MKK7-overexpression (**Fig. 5**). To circumvent the ETI triggered by HopZ1a in the Col-0 wild type background, we generated MKK7-DEX transgenic plants in the *zar1-1* knockout mutant background, in which HopZ1a does not trigger immunity (Lewis et al., 2010). Expression of HopZ1a from *Pto* DC3000 was able to suppress the attenuation of growth caused by overexpression of MKK7 (**Fig. 5a**), supporting that MKK7 is a relevant target for HopZ1a virulence activity. To determine whether bacteria-delivered HopZ1a is able to suppress MKK7-associated PTI responses, we used *Pseudomonas fluorescens* strain Pf55 (hereafter Pf55), a non-pathogenic strain expressing a heterologous functional T3SS (Jamir et al., 2004). Inoculation with Pf55 resulted in the accumulation of callose, which was noticeably higher in MKK7-overexpressing plants (**Fig. 5b**). However, HopZ1a expression from Pf55 inhibited callose deposition, and even abolished the enhanced accumulation caused by MKK7 overexpression (**Fig. 5b**), indicating that HopZ1a is sufficient to suppress callose deposition mediated by MKK7. Last, we analyzed whether HopZ1a was able to suppress MKK7-dependent PR1 accumulation. Overexpression of MKK7 has been described to induce *PR1* gene expression (Zhang et al., 2007b). Accordingly, MKK7-overexpressing plants treated with water (as mock control) or inoculated with Pf55 displayed local PR1 accumulation in the inoculated tissues (**Fig. 5c**). Noticeably, local PR1 accumulation was completely abolished in plants inoculated with Pf55 expressing the HopZ1a effector protein (**Fig. 5c**), indicating that HopZ1a is also capable of suppressing MKK7-dependent PR1 accumulation.

**FIGURE 5.**
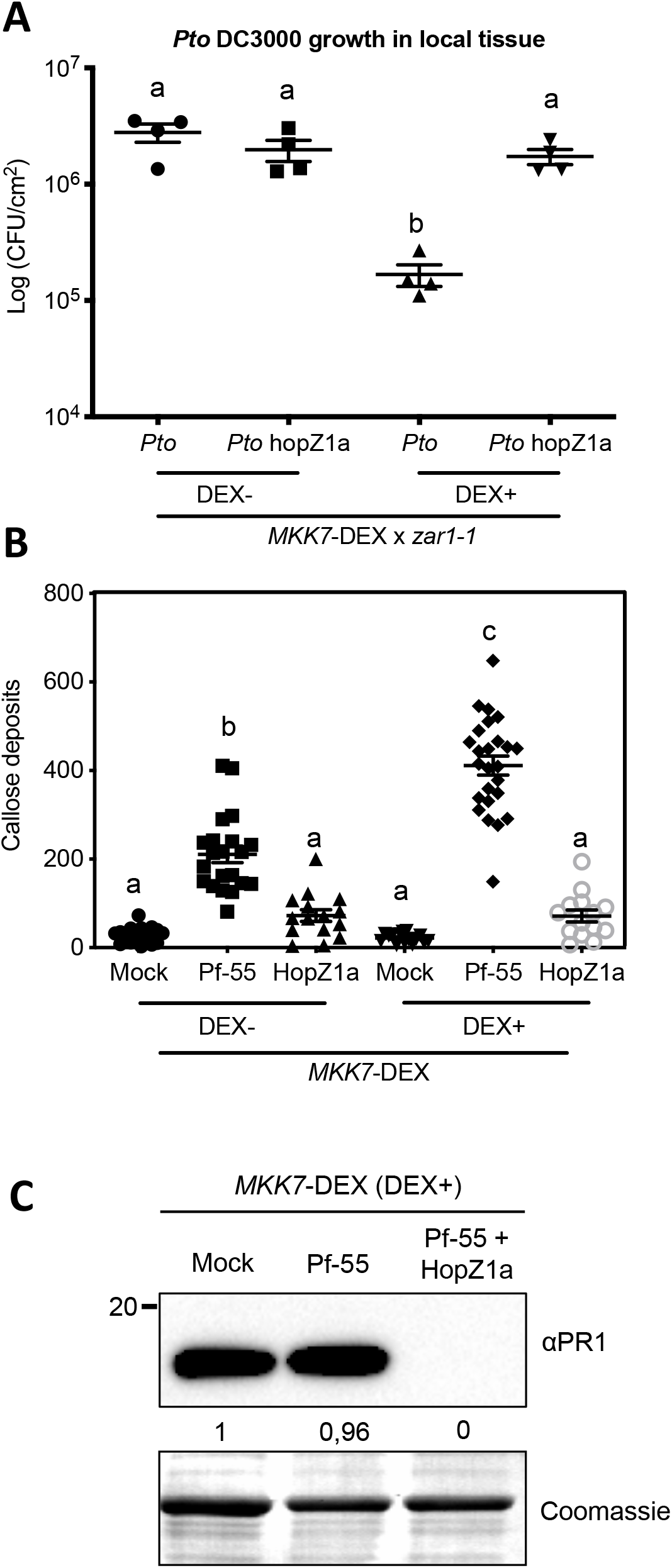
HopZ1a interferes in the MKK7 defense activation pathway. (**A**) Bacterial multiplication assay in MKK7-DEX x *zar1-1* plants. Leaves were infiltrated with either DEX (DEX+) or Mock solution (DEX−). Two hours after treatment, leaves were infiltrated with a 5 × 10^4^ cfu/ml suspension of either *Pto* DC3000 or *Pto* DC3000 expressing HopZ1a. The graphs show bacterial titers at 4 dpi. Lines represent mean values and error bars represent standard error (n=4). Individual values are shown. Statistical differences were determined using one-way ANOVA (α= 0.05) with Tukey’s multiple comparisons test and different letters indicate statistical significance. (**B**) Quantification of callose deposits in MKK7-DEX plants pretreated with either DEX (DEX+) or mock solution (DEX−). Twenty-four hours after treatment, same leaves were infiltrated with 10 mM MgCl_2_ (mock), 5 × 10^5^ cfu/ml suspension of Pf-55 (empty vector) or a 5 × 10^5^ cfu/ml suspension Pf-55 expressing HopZ1a. Eighteen hours after bacterial inoculation, leaves were stained with aniline blue and observed under UV fluorescence microscope. Lines represent average values (n=24) and error bars represent standard error. Individual values are shown. Statistical differences were determined using one-way ANOVA (α= 0.05) with Tukey’s multiple comparisons test and different letters indicate statistical significance. (**C**) PR1 levels in *MKK7*-DEX x *zar1-1* plants pretreated with either DEX (DEX+) or mock solution (DEX−). Two hours after treatment, same leaves were infiltrated with 10 mM MgCl_2_ (mock), 5 × 10^5^ cfu/ml suspension of Pf-55 (empty vector) or a 5 × 10^5^ cfu/ml suspension Pf-55 expressing HopZ1a. Samples were taken at 48 hpi. Coomassie staining is shown as loading control.

#### MKK7-dependent SAR is suppressed by HopZ1a

MKK7 has been shown to participate in SAR signaling in Arabidopsis after *Pto* DC3000 inoculation (Zhang et al., 2007b). Since HopZ1a suppresses SAR (Macho et al., 2010), we set out to determine whether HopZ1a suppresses MKK7-dependent SAR. We used Arabidopsis MKK7-DEX leaves, either (locally) treated with DEX or not, 2 hours prior to carrying out a primary (local) inoculation in these same leaves with a *P. syringae* Pf55 strain constitutively expressing HopZ1a from a stable plasmid (**Fig. 6**). Three days later, we either performed a secondary infection on distal (systemic) leaves with a fully virulent strain (*Pto* DC3000) (**Fig. 6a**), or extracted proteins to quantify systemic PR1 accumulation in distal (systemic) non-inoculated leaves (**Fig. 6b**). Systemic leaves were not treated with DEX at any point. We finally quantified bacterial multiplication on systemic, secondary infection sites as a direct and biologically relevant measurement of SAR (**Fig. 6a**). As controls, we also performed inoculation at primary sites using either infiltration buffer only (mock inoculation), or Pf55 not expressing the HopZ1a gene. While systemic multiplication of *Pto* DC3000 was significantly reduced in MKK7-DEX-induced plants that had been either mock-inoculated in primary (local) leaves, or inoculated with Pf55 (**Fig. 6a**), in plants inoculated in primary (local) leaves with HopZ1a-expressing Pf55, multiplication of *Pto* DC3000 in secondary (distal) sites was maintained up to levels similar to those reached in control non-induced MKK7-DEX plants. Further, PR1 accumulation in secondary, systemic sites of infection was suppressed only in those plants that had been previously inoculated in primary (local) leaves with the HopZ1a-expressing Pf55 strain (**Fig. 6b**). Taken together, these results indicate that HopZ1a is capable of suppressing the SAR response specifically triggered through MKK7 expression.

**FIGURE 6.**
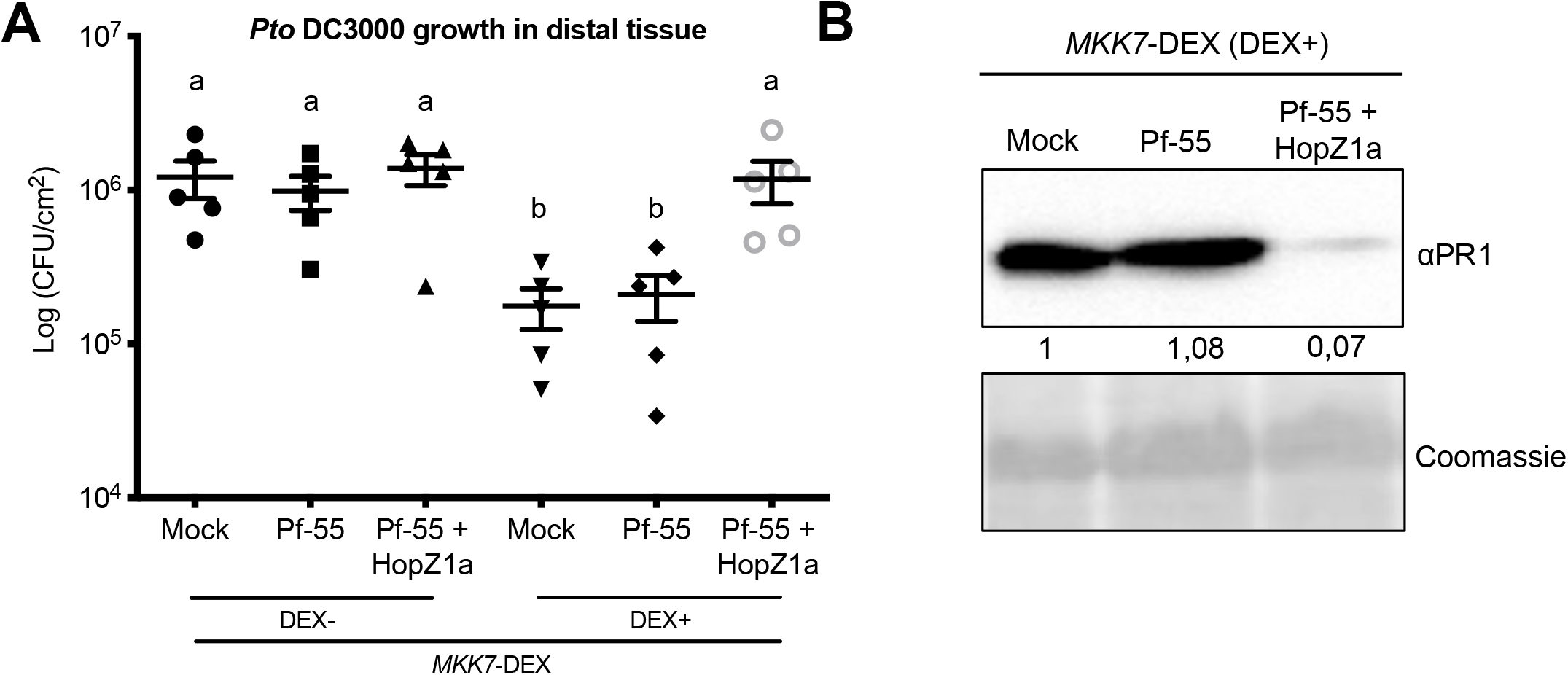
HopZ1a suppresses *AtMKK7-* triggered SAR. (**A**) Bacterial multiplication assay in distal leaves of MKK7-DEX plants pre-inoculated with the indicated strains. Leaves were first infiltrated with either DEX (DEX+) or Mock solution (DEX−). Two hours later, same leaves were infiltrated with 10 mM MgCl_2_ (mock), 5 × 10^5^ cfu/ml suspension of Pf-55 (empty vector) or a 5 × 10^5^ cfu/ml suspension of Pf-55 expressing HopZ1a. Three days after primary inoculation, distal leaves were inoculated with a 5 × 10^4^ cfu/ml suspension of *Pto* DC3000. Graph shows *Pto* DC3000 growth at 4 dpi. Lines represent average values (n=5) and error bars represent standard error. Individual values are shown. Statistical differences were determined using one-way ANOVA (α= 0.05) with Tukey’s multiple comparisons test and different letters indicate statistical significance. (**B**) PR1 accumulation in distal leaves of MKK7-DEX plants described in plants inoculated with the indicated strains. First, leaves were infiltrated with either DEX or mock solution. Two hours later, same leaves were infiltrated with 10 mM MgCl_2_ (mock), a 5 × 10^5^ cfu/ml suspension of Pf-55 (empty vector) or a 5 × 10^5^ cfu/ml suspension of Pf-55 expressing HopZ1a. Three days after infiltration, distal leaves were collected. Ten micrograms of total protein were loaded per sample, and Coomassie staining is shown as loading control. Results presented are representative of three independent experiments.

#### HopZ1a acetylates MKK7 in vitro

HopZ1a has been described to function as an acetyltransferase capable of strong autoacetylation *in vitro* (Lee et al., 2012; Ma et al., 2015), and also of transacetylation of some of its interacting partners in the plant (Lee et al., 2012; Jiang et al., 2013; Lewis et al., 2013; Lee et al., 2019; Bastedo et al., 2019).

To determine whether HopZ1a is able to acetylate MKK7, we performed a ^14^C-labelled-acetyl-coenzyme A (acetyl-CoA) transferase reaction *in vitro*, in the presence of MKK7 and either HopZ1a or HopZ1a^C216A^. As previously described, HopZ1a was strongly autoacetylated, while HopZ1a^C216A^ was not (**Fig. 7, Fig. S3**). More importantly, MKK7 was acetylated in the presence of HopZ1a, but not in the presence of HopZ1a^C216A^ (**Fig. 7, Fig. S3**), supporting the notion that HopZ1a acetylates MKK7 *in vitro* in a manner dependent on the integrity of its catalytic site. We also analyzed the acetyltransferase activity of HopZ1a^K289R^, a mutant that has been described to display reduced transacetylation activity (Ma et al., 2015). Interestingly, MKK7 was not acetylated in the presence of HopZ1a^K289R^ (**Fig. S3**), supporting the notion of HopZ1a K289 contributing to HopZ1a transacetylation activity of MKK7, as previously suggested (Cecchini *et al.*, 2015).

**FIGURE 7.**
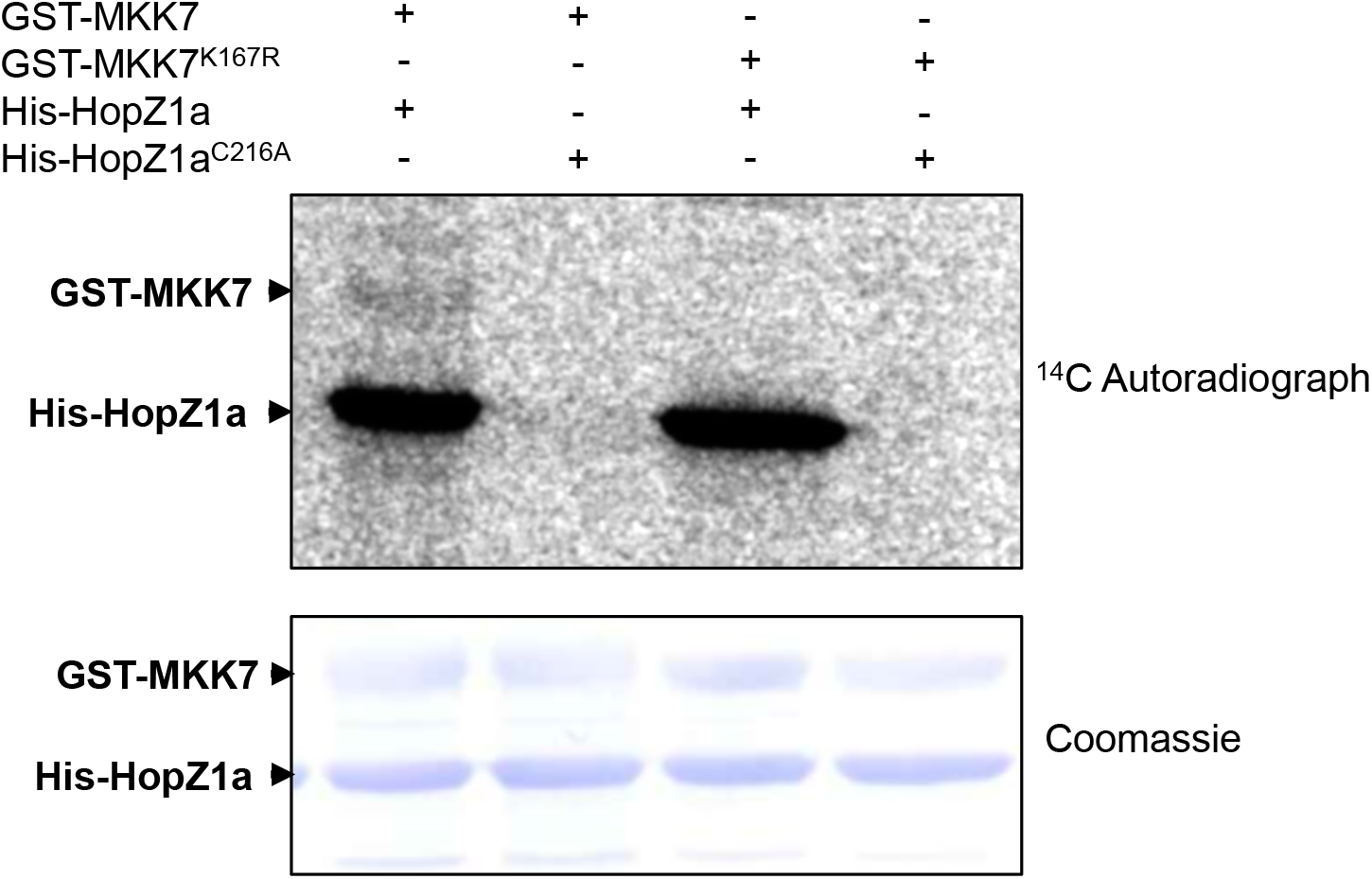
HopZ1a acetylates MKK7 *in vitro*. *In vitro* acetylation assay. Five μg of GST-MKK7 or GST-MKK7^K167R^ were incubated with 3 μg of 6xHis-HopZ1a or 6xHis-HopZ1a^C216A^ in acetylation buffer containing ^14^C-Acetyl CoA. Samples were separated by SDS-PAGE and proteins were transferred to a PVDF membrane. The membrane was exposed to an imaging plate for one week and acetylation was detected by autoradiography. Coomassie staining is shown as loading control.

HopZ1a acetylation of the Arabidopsis pseudokinase ZED1, a decoy/adaptor involved in ZAR1-dependent HopZ1a-triggered immunity, takes place on threonine residues located in positions 125 and 177 of its amino acid sequence (Lewis et al., 2013), with T177 in particular located within the catalytic loop (**Fig. S4c**). The catalytic loop constitutes a conserved domain that in active kinases includes the proton acceptor motif (HRD) essential for kinase activity (Lewis et al., 2013)**; Fig. S4)**. However, the catalytic loop of MKK7 lacks threonine residues (**Fig. S4**). Interestingly, VopA, another effector from the YopJ family, modifies a lysine residue located within the catalytic loop of its mammalian target MKKs, disrupting ATP binding and inactivating the kinase (Trosky et al., 2007). Thus, we reasoned that highly conserved lysine residue K167, located within the catalytic loop of MKK7 (**Fig. S4**) was a good candidate for HopZ1a acetylation.

To test K167 potential relevance for HopZ1a interference with MKK7 activity, we introduced a point mutation into MKK7 by substituting residue K167 with an arginine to generate mutant MKK7^K167R^ and used this mutant protein as a substrate for HopZ1a acetylation *in vitro*. Results of this assay show that mutation K167R prevents HopZ1a-mediated acetylation of MKK7 (**Fig. 7**, **Fig. S3**), suggesting that K167 is a major target residue for HopZ1a acetylation.

### Lysine residue K167 is required for full MKK7 activity *in vitro* and *in planta*

It has been previously shown that MKK7-dependent activation of plant immunity requires MKK7 kinase activity (Zhang et al., 2007b). MKK7 autophosphorylation activity is absent in the MKK7^K74R^ mutant version of the protein, in which a lysine within the ATP binding pocket has been changed to an arginine (Dai et al., 2006). To determine whether the K167 residue is necessary for MKK7 kinase activity, we carried out *in vitro* GST-MKK7, GST-MKK7^K74R^, and GST-MKK7^K167R^ autophosphorylation assays (**Fig. 8a)**. Autophosphorylation was strongly reduced in the GST-MKK7^K167R^ mutant version (**Fig. 8a**), and completely abolished in the control GST-MKK7^K74R^ mutant version. We also detected a mobility shift in SDS-PAGE gels in GST-MKK7 samples, which displayed a slower migration rate suggestive of protein phosphorylation, which was absent in both GST-MKK7^K74R^ and GST-MKK7^K167R^ samples (**Fig. 8b**). These results indicate that K167 plays a key role in MKK7 kinase activity.

**Figure 8.**
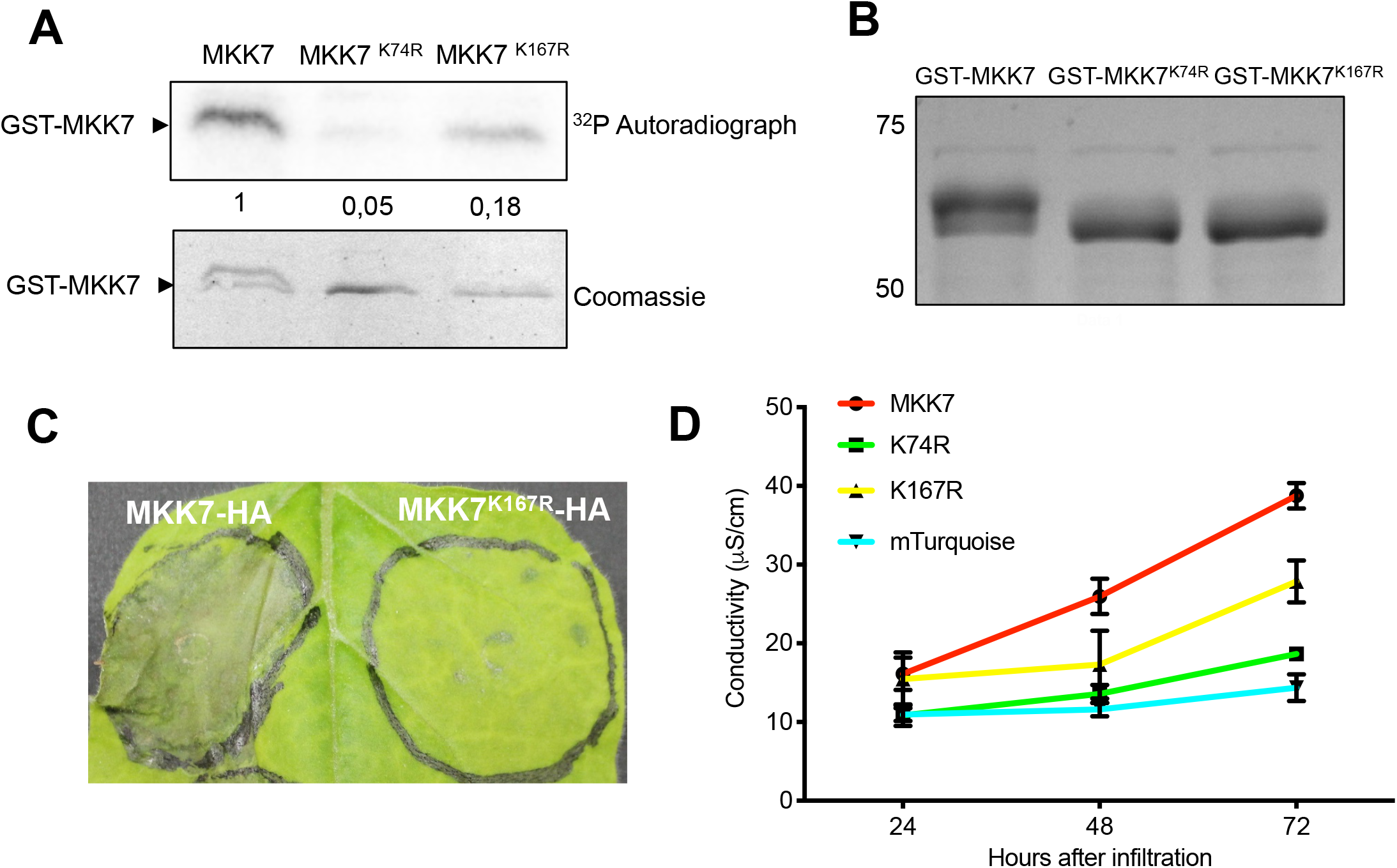
K167 is required for full MKK7 activity *in vitro* and *in planta.* (**A**) *In vitro* kinase assay. One μg of GST-MKK7, GST-MKK7^K74R^ or GST-MKK7^K167R^ were incubated in kinase buffer containing ^32^P-γ-ATP. Samples were separated by SDS-PAGE and proteins were transferred to a PVDF membrane. The membrane was exposed to an imaging plate for one day and autophosphorylation was detected by autoradiography. Coomassie staining is shown as loading control. (**B**) Five μg of GST-MKK7, GST-MKK7^K74R^ or GST-MKK7^K167R^ were separated by SDS-PAGE. SDS-PAGE were stained with Coomassie staining. (**C**) Cell death induced by MKK7. Transient expression in *N. benthamiana* leaves inoculated with *A. tumefaciens* carrying either MKK7-HA or MKK7^K167R^ –HA constructs, as indicated. Pictures were taken at 48 hpi. (**D**) Ion leakage assay in *N. benthamiana* leaves inoculated with *A. tumefaciens* carrying MKK7-HA, MKK7^K74R^ –HA or MKK7^K167R^ –HA constructs. Conductivity was measured at the indicated time points. Graph indicates mean values (n=3) and error bars indicate standard error. Experiments were repeated three times with similar results.

To confirm that K167 is also key for MKK7 activity *in planta*, we expressed MKK7 and its mutant derivatives MKK7^K167R^ and MKK7^K74R^ in *N. benthamiana* leaves under the control of a constitutive promoter using *Agrobacterium*-mediated transient expression (**Fig. 8c**). Transient overexpression of MKK7 resulted in the manifestation of macroscopic cell death in *N. benthamiana* tissues (**Fig. 8c**), most likely as a result of immune activation, as previously suggested by the failure to generate *Arabidopsis* transgenic plants expressing this kinase from a 35S promoter (Dai et al., 2006; Zhang et al., 2007b). Interestingly, neither the *bona fide* mutant protein MKK7^K74R^ nor MKK7^K167R^ expression elicited macroscopic cell death (**Fig. 8c**), nor ion leakage associated to the onset of cell death in *N. benthamiana* tissues (**Fig. 8d**). These results indicate that the K167 residue, targeted for acetylation by HopZ1a, is essential for MKK7 activity *in planta*.

## DISCUSSION

In this work we identify and characterize the interaction between the *P. syringae* T3E HopZ1a and Arabidopsis MKK7, a positive regulator of plant defense. We show that this interaction results in acetylation of MKK7, likely in a lysine residue essential for its kinase activity, and demonstrate that bacteria-delivered HopZ1a suppresses MKK7-dependent defense responses, thus providing a molecular mechanism for HopZ1a simultaneous suppression of PTI, ETI, and SAR signaling.

While we have shown that HopZ1a interference with MKK7 may explain HopZ1a-dependent defense suppression at all levels (PTI, ETI, and SAR), our results do not rule out additional HopZ1a interactions with other Arabidopsis MKKs, considering the overall conservation among this class of MAP kinases (**Fig S4**). As an example, the *P. syringae* T3E HopF2 has been described to interact with MKK4/5 and to modify MKK5, but can also interact with other Arabidopsis MKKs (Wang et al., 2010). Similarly, YopJ acetylates mammalian MKK6 (Mukherjee et al., 2006), but also MKK4 and MKK7 (Paquette et al., 2012). HopZ1a interference with other positive regulators of defense like the redundant pairs MKK1/2 or MKK4/5 (reviewed by (Meng & Zhang, 2013) could contribute to HopZ1a-associated phenotypes. In any case, only MKK7 has been proved to be essential for SAR activation to date (Zhang et al., 2007b), which implies a special role for this novel target, at least in relation to HopZ1a suppression of systemic defense (**Fig. 6**).

Our results indicate that HopZ1a acetylates MKK7, most likely in a conserved, essential lysine located in the catalytic loop of the kinase, probably involved in the coordination of ATP binding (**Fig. 7; Fig S4**). HopZ1a has been previously described to auto-acetylate in lysine residue K289 (Lee et al., 2012; Ma et al., 2015; Rufián et al., 2015). In fact, a number of effectors of the HopZ1 superfamily, like YopJ from *Yersinia*, AvrA from *Salmonella,* and VopA from *Vibrio* can also acetylate lysine residues on its corresponding target MKKs, typically resulting in inhibition of kinase activity and suppression of immune responses (Mukherjee et al., 2006; Trosky et al., 2007; Jones et al., 2008; Paquette et al., 2012). In the case of VopA, acetylation of a conserved lysine located on the catalytic loop of mammalian MKK6 disrupts ATP binding and inactivates the kinase (Trosky et al., 2007). We propose an equivalent mechanism for HopZ1a interference with MKK7 kinase activity.

However, HopZ1a might also modify additional MKK7 residues to interfere with kinase function. HopZ1a has been described to auto-acetylate in two serine residues (S349 and S351) that are required for acetyltransferase activity *in vitro*, virulence activity *in planta*, and interaction with the co-factor IP6 (Ma et al., 2015). The T3Es YopJ, AvrA, and VopA also acetylate key serine and threonine residues in the activation loop of their target host kinases (Mittal et al., 2006; Mukherjee et al., 2006; Trosky et al., 2007; Jones et al., 2008; Paquette et al., 2012; Meinzer et al., 2012; reviewed by Ma & Ma, 2016). Therefore, since ours has been a direct approach, we cannot rule out additional HopZ1a acetylations of serine and/or threonine residues on MKK7. In this sense, Popescu et al. (2009) showed that wild type MKK7 was more active than the constitutively active mutant version, where essential serine and threonine residues in the activation loop sequence were replaced with glutamic acid, suggesting the possibility that MKK7 is also regulated through the phosphorylation of other residues located within or outside the activation loop.

It is however likely that HopZ1a interference with other participants in local and systemic defense will also contribute to different HopZ1a-associated suppression phenotypes. To date, HopZ1a have been proposed to interact with a number of plant proteins, among them several kinases associated to immune signaling like PBS1-like (PBL) (Bastedo et al., 2019), or SZE1 and SZE2 (Liu et al., 2019). Current knowledge indicates that a single effector can have multiple host target proteins. In the extreme, some effectors can exert a single activity on a whole class of host target proteins without particular need for specificity (e.g. AvrPto is a kinase inhibitor that can target receptor kinases, whether they are involved in immunity or not) (as discussed by (Macho & Zipfel, 2015)). With a less generalist approach, HopZ1a seems to act on a number of different plant kinases associated to immune signaling, preferentially RLCKs and MAPKs, interfering with their function in susceptible plants or giving itself away in resistant plants, through acetylation of key residues. The structural conservation amongst different kinases likely facilitates a certain laxity in target selection. Simultaneous action on several kinase-regulated links of the same plant signaling process might result on a more substantial interference. Interestingly, the HopZ1a homolog HopZ3 also interacts with a number of plant RLCKs (*e*.*g*. RIPK, PBS1, BIK1, and PBL1) and also with MAP kinases (MPK4), acetylating at least some of these targets to interfere with plant defense (Lee et al., 2015).

The proposed scenario does not preclude a role for those additional targets, other than kinases, previously described for HopZ1a (Lee et al., 2012; Jiang et al., 2013; Albers et al., 2019). Some targets, like remorin (Albers et al., 2019), are directly associated with the aforementioned kinases, as co-participants in defence signaling. In other instances, the virulence effect described for HopZ1a interference with a particular interactor might be, in fact, indirect. This might be the case for the destruction of the plant microtubule network associated to HopZ1a interaction with tubulin (Lee et al., 2012). Lee and collaborators (2012) discussed that microtubule destruction might be an indirect effect of HopZ1a acting on a then unidentified protein and suggested a MAPK as a likely possibility. In support of this notion, a recent report has shown that HopZ1a interacts with kinesins (Lee et al., 2019), microtubule-associated molecular motors that achieve transmission of a variety of signals in close association to the MAPK cascade (reviewed by (Liang & Yang, 2019). In relation to this, MKK7 interacts with MPK6, which has been reported to localize, among other subcellular localizations, to microtubules (Shen *et al.*, 2019; Muller *et al.*, 2010), and microtubule-associated dynamin-related proteins 2A and 2B have been identified as phosphorylation substrates of the MKK7-MPK3/6 module in *Arabidopsis* (Strack *et al.*, 2013; Huck *et al.*, 2017).

HopZ1a associates through myristoylation to the plant plasma membrane (Lewis et al., 2008), where many of its interactors are located (Wilton *et al.*, 2010; Zhou *et al.*, 2014; Bastedo et al., 2019; Albers et al., 2019; Liu et al., 2019). It has been shown that under stress conditions AtMKK7 is recruited to the plasma membrane after binding to phosphatidic acid (PA) (Shen et al., 2019), a lipid secondary messenger involved in early defense signaling that targets proteins to the cell membrane (reviewed by (Zhao, 2015; Yao & Xue, 2018). Thus, HopZ1a could come in close contact with MKK7 after the latter is relocated to the plasma membrane by PA accumulation, as part of the early plant defense response to pathogen attack. This scenario will be coherent with HopZ1a ability to suppress AvrRps4, AvrRpm1 and AvrRpt2-triggered ETI (Macho et al., 2010), and also with AtMKK7 contribution to RPS2-mediated immunity (**Fig. 4**). It is tempting to speculate that HopZ1a could be part of a membrane-associated protein complex involving multiple host targets plus those partner effectors shielded by HopZ1a from triggering ETI, as has been shown for HopZ3 (Lee et al., 2015).

In this work we show that HopZ1a interference with AtMKK7 results in the suppression of SAR. This is one of the most distinctive virulence phenotypes resulting from HopZ1a-MKK7 interaction. Effective systemic immunity requires the rapid generation of inducing signals at the local site of infection, its long-distance translocation typically via the phloem, as well as signal perception in systemic tissues (reviewed by (Shine et al., 2019). A diverse network of inducers, carefully balanced by an extensive crosstalk, participates in SAR signaling, including phytohormones, primary/secondary metabolites, lipid derivatives, and proteins (Shine et al., 2019). Generally speaking, SAR is signaled *via* two branches operating in parallel, one regulated by the phytohormone salicylic acid (SA) and the second regulated in combination by azelaic acid (AzA), G3P, nitric oxide (NO), and Reactive Oxygen Species (ROS) (Klessig et al., 2018; Shine et al., 2019). Additionally, feeding into both branches of the SAR signaling pathway is pipecolic acid (Pip), a metabolite that is an indispensable switch for the activation of SAR, is transported systemically, and is synthesized from lysine in a process dependent on the aminotransferase ALD1 (ABERRANT GROWTH AND DEATH 2 (AGD2)-LIKE DEFENSE RESPONSE PROTEIN 1) (Navarova *et al.*, 2012; Bernsdorff *et al.*, 2016).

MKK7 downstream targets can provide an insight on the molecular events following HopZ1a interference with MKK7 in regard to systemic immunity suppression. In the presence of the MAPK module MKK7/MPK10 the DNA-binding factor TGA1 is phosphorylated (Popescu et al., 2009). TGA1 and closely related TGA4 positively regulate basal resistance against pathogens and, more relevantly, are also required for SAR (Kesarwani *et al.*, 2007; Shearer *et al.*, 2012; Sun *et al.*, 2018). TGA1 and TGA 4 activate the expression of ISOCHORISMATE SYNTHASE 1 (ICS1), leading to an increase in SA biosynthesis (Sun et al., 2018), thus contributing to signaling through the first of the abovementioned SAR signaling branches. Further, upon pathogen infection, TGA1 and TGA4 contribute to the induction of SYSTEMIC ACQUIRED RESISTANCE DEFICIENT 1 (SARD1) and CALMODULIN-BINDING PROTEIN 60g (CBP60g) that in turn, by activating ALD1 and SARD4 expression, promote the biosynthesis of Pip (Navarova et al., 2012; Bernsdorff et al., 2016; Sun et al., 2018), thus contributing to both SAR signaling branches. MKK7 also phosphorylates MPK3 and MPK6, and targets downstream this module have been associated to plant defense (Yoo *et al.*, 2008; Jia *et al.*, 2016; Huck et al., 2017; Shen et al., 2019). Interestingly, MPK3 interacts with AZELAIC ACID (AZA) INDUCED 1 (AZI1), a lipid-transfer protein that mediate AzA uptake and mobilization on the SA-independent SAR signaling branch (Jung *et al.*, 2009; Pitzschke *et al.*, 2014; Cecchini et al., 2015).

In sum, positive regulation of TGA1 by the MKK7/MPK10 module is likely to contribute to MKK7-dependent SAR activation via both SAR signaling branches, by (*i*) promoting an ICS1-dependent increase in SA biosynthesis and (*ii*) promoting pipecolic acid (Pip) biosynthesis. Additionally, the MKK7/MPK3 module might also participate in SAR signaling, considering MPK3 requirement for SAR and its interaction with AZI1. Taken together, these molecular mechanisms provide a rationale for MKK7-dependent SAR activation and consequently for HopZ1a-dependent suppression of this defense response.

Further, since plant hormone immune signaling integrates a complex network in which the SA, JA, ET, or auxin crosstalk extensively, HopZ1a suppression of SAR could also derive from indirect alterations of hormone signaling. MKK7 establishes a crosstalk point between auxin signaling and the response to biotic and abiotic stresses (Zhang et al., 2008; Jia et al., 2016; Dory *et al.*, 2018). Auxin signaling, including biosynthesis, perception, localization, and transport, participates in the establishment and maintenance of SAR triggered by virulent *P. syringae* (Truman *et al.*, 2010; Bennett *et al.*, 1996; Kepinski & Leyser, 2005; Dharmasiri *et al.*, 2006). Jasmonates also have been proposed to participate in SAR signaling (Truman *et al.*, 2007; Chaturvedi *et al.*, 2008), and HopZ1a interacts in Arabidopsis with JASMONATE ZIM DOMAIN (JAZ) transcriptional repressors, inducing its degradation and thus promoting JA-responsive gene expression (Jiang et al., 2013), raising the possibility that its interference with JA signaling also contributes to HopZ1-mediated suppression of SAR. Further studies will be necessary to understand the integrated impact of HopZ1a virulence activities in plant hormone and defense signaling.

## Acknowledgements

We wish to thank Dr. Zhonglin Mou (University of Florida) for his kind gift of Arabidopsis *MKK7*-DEX and *asMKK7* transgenic plants, and Alicia Esteban del Valle for help with the *zar1-1* crossings. We also wish to thank Dr. Vitor Amorim-Silva for technical advice, and Dr. Ainhoa Lucía, Dr. Araceli Castillo, Blanca Sabarit, and P. García Vallejo for technical help.

---

**Figure S1.**
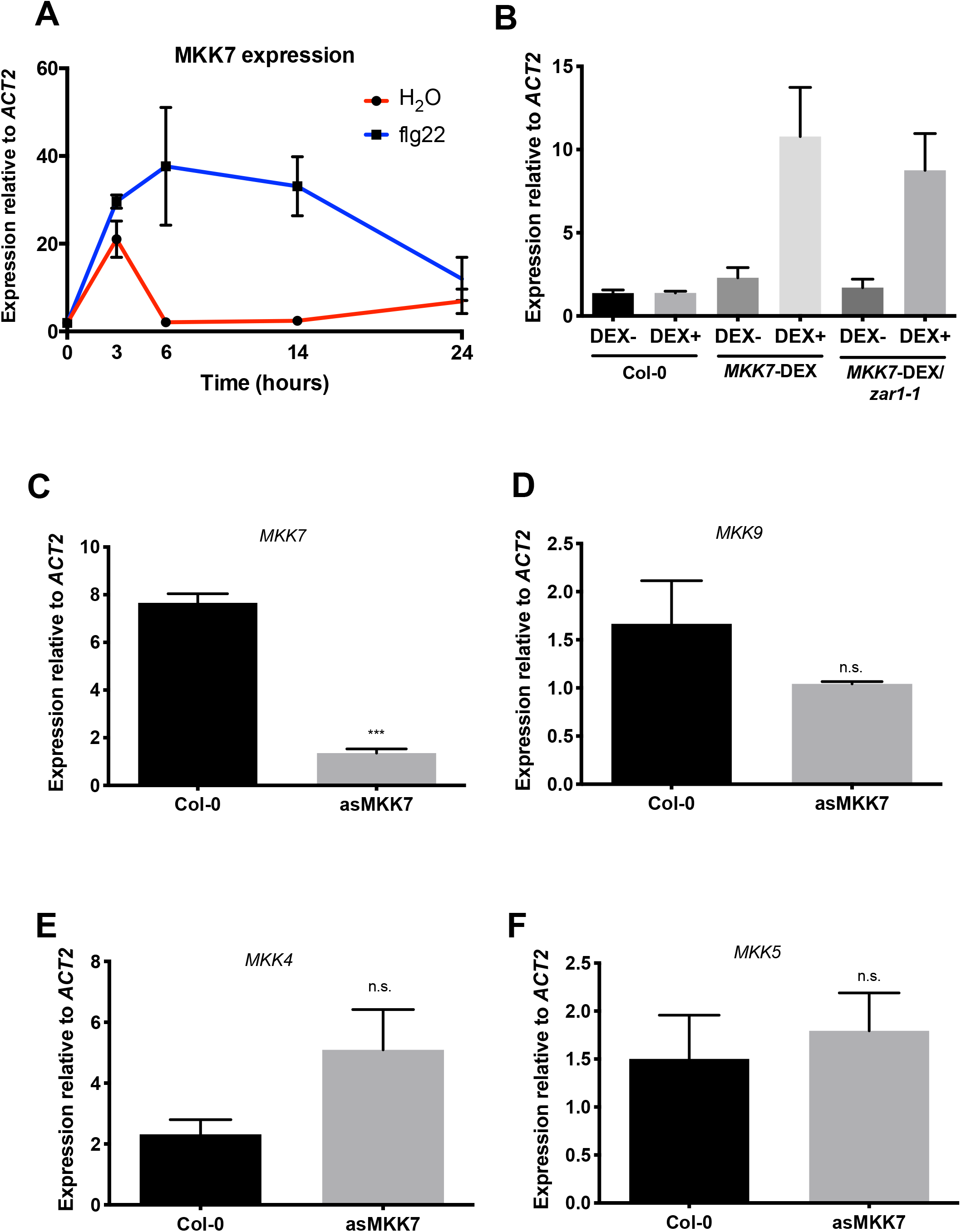
Analysis of MKK7 expression by RT-*q*PCR in Col-0 and MKK7 transgenic plants used in this work. (**A**) Relative expression of *MKK7* in Col-0 plants at different time points after treatment with 100 nM flg22 or water solution. (**B**) Relative expression of *MKK7* in Col-0, MKK7-DEX or MKK7-DEX x *zar1-1* plants after treatment with DEX or mock solution. (**C, D, E, F**) Graphs show relative expression of *MKK7*, *MKK9*, *MKK4* and *MKK5* mRNA in asMKK7 plants compared to Col-0 plants. In each experiment, error bars correspond to standard error, asterisks indicate significant differences as established by a Student’s t-test (P<0.05), while non-significant differences are marked as n.s. At*ACT2* was used as housekeeping gene and relative expression was calculated using the 2^−ΔΔCt^ method. Experiments were repeated three times with similar results.

**Figure S2.**
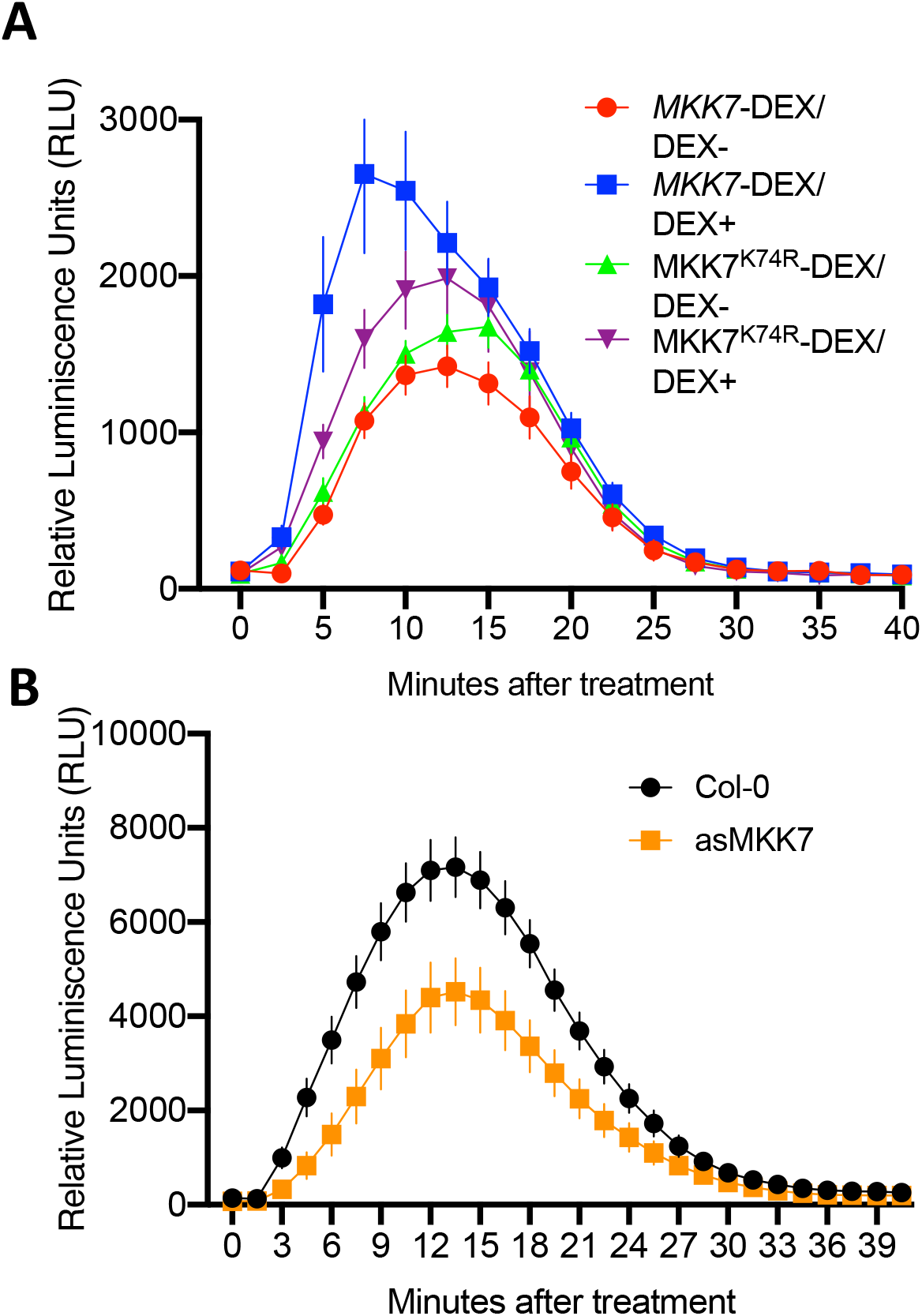
Dynamics of flg22-induced ROS burst. ROS production dynamics corresponding to the accumulated values shown in graphs displayed in Figure 2a and 2b. ROS production after treatment with 100 nM flg22 of (**A**) MKK7-DEX and MKK7^K74R^-DEX plants or (**B**) Col-0 and asMKK7 plants, measured in a luminol-based assay and represented as relative luminescence units (RLU). MKK7-DEX and MKK7^K74R^-DEX plants were treated either with a DEX or mock solution 24 hours before flg22 treatment. Error bars represent standard error (n=20). Experiments were repeated three times with similar results.

**Figure S3.**
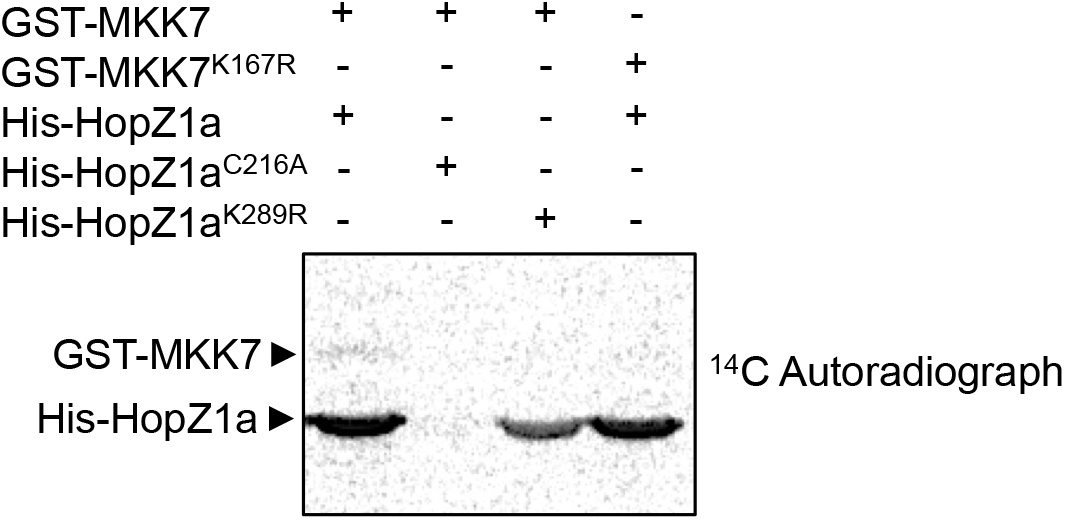
HopZ1a acetylates MKK7 in lysine 167 *in vitro*. *In vitro* acetylation assay using ^14^C-Acetyl CoA. Five μg of GST-tagged proteins (MKK7 or MKK7^K167R^) were incubated with 3 μg of 6xHis-tagged proteins (HopZ1a, HopZ1a^C216A^ or HopZ1a^K289R^) in acetylation buffer containing ^14^C-Acetyl CoA. Samples were separated by SDS-PAGE and proteins were transferred to a PVDF membrane. The membrane was exposed to an imaging plate for one week and acetylation was detected by autoradiography.

**Figure S4.**
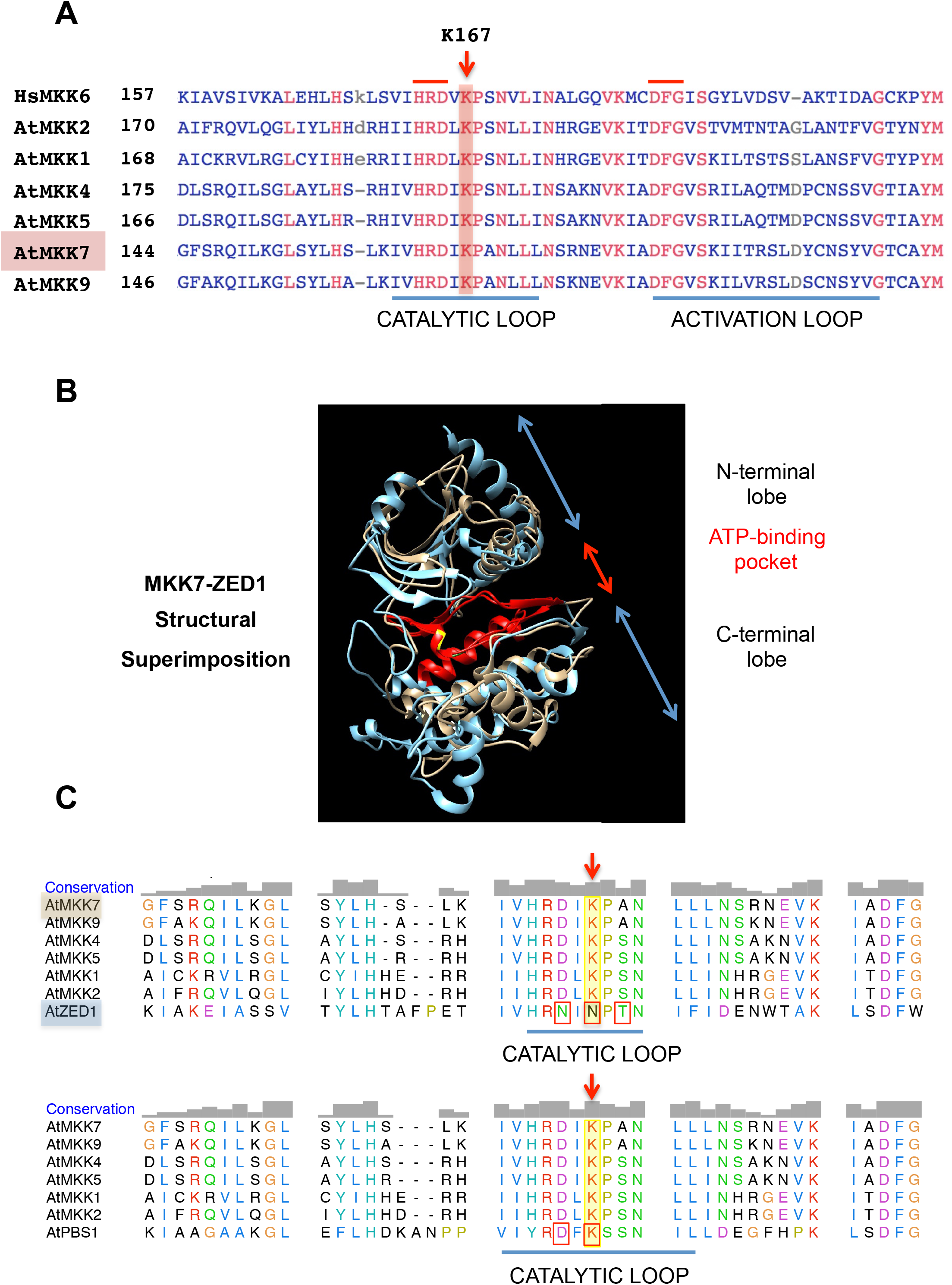
Arabidopsis MKK7 amino acid sequence and structure comparison with relevant kinases. (**A**) Multiple sequence alignment of *Arabidopsis thaliana* MAP kinase kinases AtMKK7 (NP_173271.1), AtMKK9 (NP_177492.1), AtMKK1 (NP_849446.1), AtMKK2 (NP_194710.1), AtMKK4 (NP_175577.1), and AtMKK5 (NP_001319606), with mammalian HsMKK6 (NP_002749.2) included as a reference. The alignment was made using the constrained-based multiple alignment tool COBALT (**Papadopoulos and Agarwala, 2007**) available at the National Center for Biotechnology Information (NCBI). The displayed sequence encompasses the kinases catalytic and activation loops (underlined) and surrounding regions; the first amino acid position for each sequence is indicated (left). Boundaries of the catalytic loop are determined by the position of the β6 and β7 strands (**Fabbro et al. 2015**; **Nolen et al. 2004**) predicted for AtMKK7 using the protein modelling portal PHYRE2 (**Kelley et al. 2015**). The essential HRD (catalytic) and DGF motifs are indicated with red lines over the first aligned sequence. Amino acid positions in red are conserved in all aligned sequences. The essential, conserved lysine residue of the catalytic loop acetylated by HopZ1a (K167) is indicated by a red arrow. (**B**) Image shows a superimposition of AtMKK7 and AtZED1 protein structures obtained using the program UCSF Chimera (**Pettersen et al. 2004**), illustrating the common overall fold of all eukaryotic protein kinases. For AtMKK7 (gold) we used a protein model obtained using the protein modelling server iTASSER (**Yang and Zhang 2015**), and for AtZED1 (light blue) a homology model from the SWISS-MODEL repository (**Bienert et al. 2017**). The structure corresponding to the ATP binding pocket (colored in red) is seen as a cleft between the amino- and carboxy-terminal lobes of the kinases. The conserved lysine residue (K167) of the catalytic loop that is acetylated in AtMKK7 by HopZ1a is colored in yellow. (**C**) Multiple alignments obtained using the program UCSF Chimera, based on linear sequence and structural constraints (**Pettersen et al. 2004**), including AtMKK7 (Q9LPQ3), AtMKK9 (Q9FX43), AtMKK1 (Q94A06), AtMKK2 (Q9S7U9), AtMKK4 (O80397), AtMKK5 (Q8RXG3), and HopZ1a-interacting pseudokinase AtZED1 (Q8LGB6) (above) or active kinase AtPBS1 (Q9FE20) (below). Displayed on top of the alignment (grey columns) is a graphical representation of the RMSD (root-mean-square deviation of atomic positions) estimating the average distance between the backbone atoms of the superimposed proteins, used here as a quantitative measure of similarity between the two protein structures. The sequences displayed encompass the catalytic loop (underlined) and surrounding regions. Boundaries of the catalytic loop are those predicted for AtZED1 (**Lewis et al. 2013**), or for AtPBS1 subdomain VIb (**Swiderski and Innes. 2001**). Relevant residues for AtZED1 (above) are boxed in red, including asparagine N173 that substitutes the proton-acceptor aspartic acid essential for catalysis in active kinases, and threonine T177 that is acetylated by HopZ1a and is key for ZED1-PBS1 interaction (**Lewis et al. 2013**). Relevant residues for AtPBS1 (below) are also boxed in red, including the proton-acceptor aspartic acid essential for catalysis (D213), and the lysine residue (K215) equivalent to the lysine residue acetylated in AtMKK7 1a (K167, indicated by the red arrow). In AtZED1, the lysine residue in all AtMKKs and in AtPBS1 is substituted by N175 (also boxed)

